# Microglia maintain homeostatic conditions in the developing rostral migratory stream

**DOI:** 10.1101/2022.05.17.492317

**Authors:** Sarah J. Meller, Lexie Hernandez, Eduardo Martin-Lopez, Teresa Liberia, Charles A. Greer

**Author notes:** Address correspondence to Dr. Charles A. Greer. Author contributions: SJM designed research, performed research, analyzed data, and wrote the paper. LH performed research. EML performed research. TL contributed analytic tools. CAG designed research and reviewed the paper.

## Abstract

Microglia invade the neuroblast migratory corridor of the rostral migratory stream (RMS) early in development. This work examines how microglia maintain the homeostatic conditions permissive to neuroblast migration in the RMS during the early postnatal period. GFP labeled microglia in *CX3CR-1^GFP/+^* mice assemble primarily along the outer borders of the RMS during the first postnatal week, where they exhibit predominantly an ameboid morphology and associate with migrating neuroblasts. Microglia ablation for 3 days postnatally does not impact the density of pulse labeled BrdU+ neuroblasts nor the distance migrated by tdTomato electroporated neuroblasts in the RMS. However, microglia wrap DsRed-labeled neuroblasts in the RMS of P7 *CX3CR-1^GFP/+^;DCX^DsRed/+^* mice and express the phagocytic markers CD68, CLEC7A and MERTK, suggesting active phagocytosis of neuroblasts in the developing RMS. Microglia depletion for 14 days postnatally further induced an accumulation of DCX+ neuroblasts and CC3+ apoptotic cells in the RMS, a wider RMS and extended patency of the lateral ventricle extension in the olfactory bulb. These findings illustrate the importance of microglia phagocytosis in maintaining the homeostasis of the early postnatal RMS.

**SIGNIFICANCE STATEMENT:** Microglia are brain-resident immune cells responsible for both maintaining homeostatic conditions necessary for normal neurodevelopment as well as orchestrating the brain’s response to environmental insults. The effects of microglia-mediated immune response during development may be of special relevance to the olfactory system, which is unique in both its vulnerability to environmental insults as well as its extended period of neurogenesis and neuronal migration. The work presented here examines how microglia maintain homeostatic conditions in the neuroblast migratory corridor of the rostral migratory stream (RMS) in the olfactory system during early postnatal development. Our findings illustrate the importance of microglia phagocytosis in the early postnatal RMS and provides insights into microglia function during periods of neurogenesis and neuronal migration.

## INTRODUCTION

Microglia are brain-resident immune cells responsible for both maintaining the homeostatic conditions necessary for normal neurodevelopment as well as orchestrating the brain’s response to environmental insults (Pierre et al., 2017; Tay et al., 2017; Hammond et al., 2018; Lenz and Nelson, 2018; Li and Barres, 2018; Ferro et al., 2021). The importance of microglia in mediating normal neurodevelopment was recently underscored by the case of a child born without microglia. This individual carried a homozygous mutation in colony stimulating factor 1 receptor (CSF1R), which is necessary to maintain a microglia population (Nandi et al., 2012). This child presented with structural brain malformations, including: agenesis of the corpus callosum, pontocerebellar hypoplasia, cerebellar vermis hypoplasia/atrophy, a large posterior fossa, ventriculomegaly, and heterotopias (Oosterhof et al., 2019). These mirrored the findings shown by a CSF1R knockout mouse model which also demonstrated ventricular enlargement, disrupted olfactory bulb architecture and olfactory deficits (Erblich et al., 2011). Microglia functions may be of special relevance to the olfactory system, which is unique in both its vulnerability to environmental insults as well as an extended period of neurogenesis and neuronal migration.

Microglia phagocytosis is critical for maintaining homeostasis in the developing brain. Defective, necrotic and apoptotic cells all need to be eliminated by microglia phagocytosis. Microglia further adapted their phagocytic targets to include neural stem cells, synapses, axonal and myelin debris, and proteins with a high turnover rate (such as amyloid beta protein) (Prinz et al., 2017). Microglia phagocytosis is essential during critical periods of development for clearing the results of over-exuberant neurogenesis (Cunningham et al., 2013; Barger et al., 2019) and synaptogenesis (Tremblay et al., 2010; Paolicelli et al., 2011; Gunner et al., 2019; Mallya et al., 2019; Scott-Hewitt et al., 2020).

Recent studies identified a specialized microglia subtype that exhibit an ameboid morphology and phagocytic markers, including lysosomal-associated membrane protein 1 (Lamp1), CD68, and CLEC7A, in developing white matter tracts during the first postnatal week (Hagemeyer et al., 2017; Hammond et al., 2019; Li et al., 2019). These cells are not only involved in phagocytosis in developing white matter tracts (Li et al., 2019a), but also express the trophic factors IGF1 and Spp1 (Hagemeyer et al., 2017; Hammond et al., 2019; Li et al., 2019) and are involved in supporting myelination (Hagemeyer et al., 2017; Wlodarczyk et al., 2017) and projecting axons (Ueno et al., 2013). Intriguingly, this cluster of postnatal microglia that specifically expresses the phacotyic marker CLEC7A is also seen in the rostral migratory stream (RMS) in the olfactory system (Li et al., 2019), suggesting that a similar microglia subset may be involved in phagocytosis or trophic support in the developing RMS.

The RMS is an important neuroblast migratory corridor in the olfactory system. In mice, neuroblasts are continuously generated in the subventricular zone (SVZ) and migrate in chains down the RMS to the olfactory bulb (OB) (Lois and Alvarez-Buylla, 1994; Ponti et al., 2013; Bordiuk et al., 2014). Upon reaching their destination in the OB, these neuroblasts differentiate into periglomerular and granule cell interneurons and integrate into pre-existing circuits (Lledo and Saghatelyan, 2005; Whitman and Greer, 2009). Neuronal migration via the RMS is also active in the human fetus and continues for several months after birth (Sanai et al., 2011; Wang et al., 2011). The blood vessel scaffold and astrocyte tube that wraps the RMS and supports migrating neuroblasts (Bovetti et al., 2007; Snapyan et al., 2009; Whitman et al., 2009; Grade et al., 2013; Gengatharan et al., 2016; Fujioka et al., 2017; Todd et al., 2017) does not assemble until several weeks after birth (Law et al., 1999; Peretto et al., 2005; Nie et al., 2010; Bozoyan et al., 2012), leaving a functional void that may be filled by microglia during this period.

We therefore investigated if microglia maintain the homeostasis of the RMS and support migrating neuroblasts within it during the early postnatal period. We examined the distribution and morphology of microglia in the RMS and employed microglia ablation strategies to gain insights into their function. We found that ameboid microglia distribute along the borders of the early postnatal RMS prior to the formation of the blood vessel scaffold and astrocyte tube. Microglia ablation did not impact the migratory capacity of neuroblasts but did compromise the homeostasis of the RMS, causing an accumulation of neuroblasts and apoptotic cells that broadened the domain of the early postnatal RMS. Microglia phagocytosis therefore appears critical for maintaining the permissive environment in the RMS that enables efficient neuroblast migration.

## METHODS

### Animals

Experiments investigating microglia morphology and distribution were conducted with B6.129P2(Cg)-Cx3cr1^tm1Litt^/J (Jackson stock No: 005582) mice, which express EGFP in microglia under control of the *Cx3cr1* locus (Jung et al., 2000). The mice were maintained as homozygous colonies and crossed with C57BL/6J mice to produce heterozygous *CX3CR1^GFP/+^* mice for experiments. To examine the interactions between microglia and neuroblasts, *CX3CR-1^GFP/GFP^* mice were crossed with C57BL/6J-Tg(Dcx-DsRed)14Qlu/J (Jackson stock No: 009655), which express the red fluorescent protein variant DsRed under the control of the doublecortin (*Dcx*) locus (Wang et al., 2007). This allowed the generation of *CX3CR-1^GFP/+^;DCX^DsRed/+^* transgenic mice.

To selectively deplete microglia *in vivo,* B6.129P2(Cg)-*Cx3cr1^tm2.1(cre/ERT2)Litt/^* WganJ (Jackson stock No: 021160), which express a Cre-ERT2 fusion protein under the *CX3CR1* promoter (Parkhurst et al., 2013), were crossed with B6.129S6(Cg)-*Gt(ROSA)26Sor^tm1(DTA)Jpmb^*/J (Jackson stock No: 032087), which express a *loxP*-flocked STOP cassette prior to the diptheria toxin fragment A (DTA) (Ivanova et al., 2005). Homozygous *Cx3cr1^CreER/CreER^* were crossed with heterozygous *ROSA26^eGFP-DTA/+^* mice to create *Cx3cr1^CreER/+^; ROSA26^eGFP-DTA/+^* and *Cx3cr1^CreER/+^; ROSA26^+/+^* control littermates. Tamoxifen injection induces the expression of the toxic DTA subunit in microglia in *Cx3cr1^CreER/+^; ROSA26^eGFP-DTA/+^* mice.

All experiments randomly included both male and female mice, although sex comparisons were not pursued. Mice were housed with a 12 h light/dark cycle with access to standard food and water *ad libitum*. All animal care and use were performed in accordance with the [Author University] animal care committee’s regulations.

### Electroporation

To label individual migrating neuroblasts, stem cells lining the lateral wall of the lateral ventricle were electroporated with a pCAG-tdTomato at P0/P1 and killed at P2, P4, P7, P14. The pCAG-tdTomato plasmid used for electroporation was a gift from Angelique Bordey (Addgene plasmid # 83029; http://n2t.net/addgene:83029), which was maintained in E. coli strains stored at −80°C as 10% glycerol:LB stock. P0 and P1 mice were anesthetized with hypothermia prior to injection with tdTomato plasmid. The plasmid solution was injected into one of the brain lateral ventricles using a Picospritzer (General Valve Corporation). Electroporations were made by delivering 5 pulses of 35 V using a pair of gold tweezers (Genepaddles-542, Harvard Apparatus) connected to an ECM 830 electroporator (BTX Harvard Apparatus).

### Liposome Injection

Microglia were depleted in perinatal mice by the injection of clodronate-filled liposomes (Clodrosome) at P0 or P1 (Fig. 1). Injection of clodronate-filled liposomes and their subsequent phagocytosis by microglia induces cell death, reflected by a dramatic decrease in the number of microglia three days later (Cunningham et al., 2013). 2 µl of Clodrosomes or the control PBS-filled liposomes (Encapsome) were injected into the cerebral lateral ventricles of CX3CR-1^GFP/+^ mice bilaterally using a Picospritzer (General Valve Corporation) (Table 1). The effect of microglia knockdown was evaluated three days later by immunohistochemistry and confocal microscopy. To demonstrate the phagocytic capacity of microglia throughout the RMS, fluorescent liposomes containing the fluorescent dye Dil (Fluoroliposome) were injected into the cerebral lateral ventricles of P1 CX3CR-1^GFP/+^ mice and killed at P4 (Table 1). P0 and P1 postnatal mice were anesthetized with hypothermia prior to injection.

**Figure 1.**
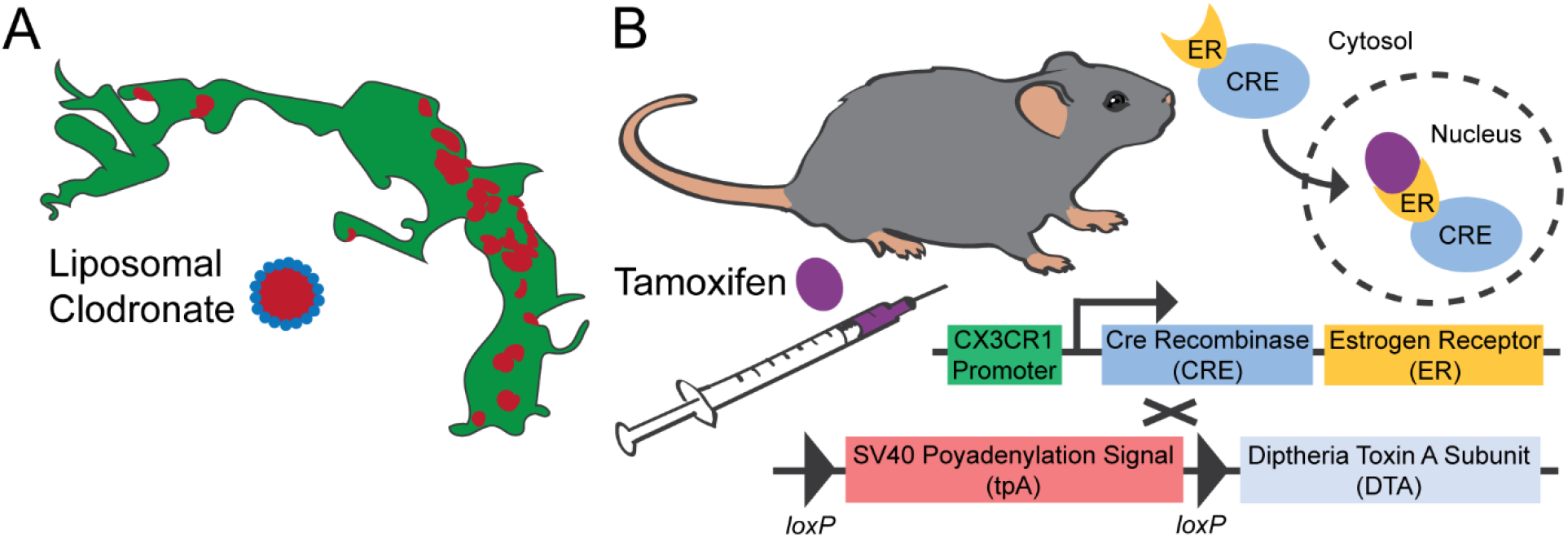
Microglia ablation methods. (A) Clodronate is encapsulated in multilamellar liposomes. When injected into the cerebral lateral ventricles, microglia phagocytose the liposomal clodronate. Clodronate is then released into the cytosol of the microglial cell, where it inhibits the mitochondrial ADP/ATP translocase and triggers apoptosis. (B) Homozygous *Cx3cr1^CreER/CreER^*, which express a Cre-ERT2 fusion protein under the *CX3CR1* promoter, were crossed with heterozygous *ROSA26^eGFP-DTA/+^* mice, which express a *loxP*-flocked STOP cassette prior to the diptheria toxin fragment A (DTA). Tamoxifen injection induces the translocation of the Cre-ERT2 fusion protein into the nucleus of microglia, excision of the *loxP*-flocked STOP cassette and expression of the toxic DTA subunit.

**Table 1.** Liposomes used in the study.

| Liposomes | Company (catalog #) |
| --- | --- |
| Standard Macrophage Depletion Kit<br>(Clodrosome® + Encapsome®) | Encapsula NanoSciences (CLD-8901) |
| Fluoroliposome®-Dil | Encapsula NanoSciences (CLD-8911) |

### BrdU Injections

To analyze the density of neuroblasts migrating in the RMS, *CX3CR-1^GFP/+^;DCX^DsRed/+^* mice were injected at P0 with 25 mg/kg of the thymidine analog BrdU (BD Pharmingen) to label newborn neuroblasts and killed at P3. To investigate the number of neuroblasts that reach the glomerular layer following microglia depletion, *CX3CR-1^GFP/+^* mice were injected with 50 mg/kg of BrdU at P1 and killed at P4.

### Tamoxifen Injections

To deplete microglia in *Cx3cr1^CreER/+^; ROSA26^eGFP-DTA/+^* mice, 30 ug/g of tamoxifen (TMX) (Sigma T5648) was injected at P0. TMX induces Cre-mediated recombination at loxP sites and removal of the terminal STOP codon sequence, enabling expression of the DTA fragment (Fig. 1). Mice were either killed at P3 or injected with TMX every subsequent three days to maintain microglia depletion before P14. *Cx3cr1^CreER/+^; ROSA26^+/+^* littermates served as controls. Microglia depletion was not successful in one of the P14 *Cx3cr1^CreER/+^; ROSA26^eGFP-DTA/+^* animals due to faulty TMX injection; this animal was excluded from analysis, as we were examining the effect of microglia knockdown rather than the effect of the TMX injection paradigm.

### Tissue Processing

For immunofluorescence, mice were deeply anesthetized with an overdose of Euthasol (Virbac) and perfused transcardially in 0.1 M PBS, pH 7.4 with 1 unit/ml heparin, followed by 4% paraformaldehyde (JT Baker) in PBS. Animals were decapitated and the brains carefully removed from the skull. The brains were postfixed in 4% PFA at 4⁰C overnight. Following PBS washes, brains were transferred to a 30% sucrose and kept at 4⁰C for 3-4 days for cryoprotection. Brains were then embedded in OCT (optimal cutting temperature) compound (Thermo Fisher Scientific). 25 µm sections were serially collected in both coronal and sagittal planes using a Reichert Frigocut Cryostat E-2800. Sections were frozen at −20⁰C until use.

For electron microscopy, the mice were perfused with 4% paraformaldehyde and 0.75% glutaraldehyde, followed by postfixation in the same fixative for 4 hours. Brains were rinsed in PBS overnight and cut on a vibratome (50 μm).

### Immunohistochemistry

Sections were thawed at 37⁰C and treated for antigen retrieval with 0.01 M citrate buffer at pH 6 and 68⁰C for 30 min. They were incubated in a blocking solution of PBS supplemented with 0.1% Triton X-100 (Sigma-Aldrich) and 0.2% bovine serum albumin (BSA) (Sigma-Aldrich) for 1 h at room temperature. Sections were incubated in a mixture of primary antibodies diluted in blocking solution at 4⁰C overnight, followed by incubation with secondary antibodies diluted in blocking solution at 1:400 for 2 h at room temperature (Table 2). Nuclei were counterstained by incubating the sections with 1 µg/ml DAPI for 10 min (Invitrogen). Sections were washed with PBS after each step. The sections finally mounted with Mowiol 4-88 (Sigma-Aldrich).

**Table 2.** Primary and secondary antibodies used in the study.

| Primary Antibodies |  |  |  |  |
| --- | --- | --- | --- | --- |
| Anti | Host | Isotype | Dilution | Company (catalog #) |
| GFP | Chicken | Polyclonal | 1:500 | Abcam (Ab13970) |
| GFAP | Mouse | Monoclonal IgG1 | 1:400 | Sigma-Aldrich (G3893) |
| CD31 | Mouse | Monoclonal IgG1 | 1:50 | Abcam (ab9498) |
| Iba1 | Rabbit | Polyclonal | 1:200 | Wako (019-19741) |
| CC3 | Rabbit | Polyclonal | 1:400 | Cell Signaling (9661) |
| MERTK | Rat | Monoclonal IgG2a | 1:100 | Invitrogen (14-5751-82) |
| CLEC7A | Rat | Monoclonal IgG2a | 1:30 | Invivogen (mabg-mdect) |
| CD68 | Rat | Monoclonal IgG2a | 1:200 | Biorad (FA-11) |
| DCX | Guinea Pig | Polyclonal | 1:400 | Sigma-Aldrich (AB2253) |

| Secondary Antibodies |  |  |  |  |
| --- | --- | --- | --- | --- |
| Anti | Host | Isotype | Dilution | Company |
| Chicken | Donkey | A488 | 1:400 | Jackson |
| Chicken | Goat | Biotin | 1:400 | Abcam |
| Mouse | Goat | A647 | 1:400 | Molecular Probes |
| Rabbit | Donkey | A488 | 1:400 | Molecular Probes |
| Rabbit | Donkey | A555 | 1:400 | Molecular Probes |
| Rat | Donkey | A647 | 1:400 | Jackson |
| Guinea Pig | Goat | A555 | 1:400 | Molecular Probes |

### Electron Microscopy

Tissue was incubated with 2% BSA (Sigma) in PBS for 30 min to block nonspecific binding sites. The tissue was incubated in a chicken polyclonal anti-GFP antibody (Abcam) in blocking buffer for 4 days at room temperature. Tissue was then incubated in biotin-conjugated goat anti-chicken IgY secondary antibodies for 1 hr at room temperature. Sections were incubated with the ABC reagent (Vector) for 1 hr, followed by a DAB peroxidase reaction. Sections were immediately processed for electron microscopy.

Stained tissue was post-fixed with 4% osmium tetroxide for 1 hr, dehydrated through graded alcohols, and polymerized in Epon between glass slides and coverslips coated previously with Liquid Release Agent (Electron Microscopy Sciences, Fort Washington, PA). Areas of interest were selected based on the DAB labeling and re-embedded in Epon. These re-embedded sections were cut using an ultramicrotome in 70 nm thick ultrathin sections. These 70 nm sections were examined with a JEOL transmission electron microscope and photographed at primary magnifications of 3,000–4,000X.

### Confocal Imaging and Processing

All images were taken using a Zeiss LSM800 confocal with either a 10X or 40X objective and images processed with Fiji (ImageJ) software. When creating tiled images of entire sagittal sections for illustrative purposes, images were taken with the 10x objective at 27 µm z-stacks (4 slices at 6.75 µm step distance.) After creating a maximum projection of each channel, a “rolling ball” algorithm was applied to correct for uneven illumination.

Cell density analysis was performed on confocal images obtained with the 10X objective. Z-stacks were 27 µm in depth (4 slices at 6.75 µm step distance.) The rostral migratory stream (RMS) was first isolated on a 16-bit maximum projection of a 638.9 x 638.9 µm image of DAPI nuclear staining with a freehand drawing tool. The area of this drawn region of interest was measured later to determine the density of stained cells and area ratio of stained processes. The ROI was drawn on the maximum projection of the DAPI channel to prevent bias for incorporating stained cells of interest. A mask of the drawn ROI was then applied to the corresponding channels stained with antibodies against GFP, GFAP, Iba1, BrdU and CC3 to quantify stained objects within this ROI.

There was increased noise in the GFP and DsRed channels, and thus we applied an additional “despeckle” function, which removes salt-and-pepper noise. Salt-and-pepper noise reduction was achieved in the CC3 channel with a “median” filter of pixel radius 1. BrdU staining was punctate, so to blur punctate staining and isolate stained nuclei a “gaussian blur” function with a pixel sigma of 2 was applied to the BrdU max projection. To segment stained pixels from background we used the Fiji automatic Otsu threshold function, except for segmenting Iba1 and CC3 stained objects, for which we used the Fiji automatic Triangle threshold function. A gray scale attribute filter with an opening function of area minimum of 100 pixels and connectivity of 8 was further applied using the MorphoLibJ integrated library and plugin (Legland et al., 2016) to isolate the cell bodies of GFP- and Iba1-stained microglia and BrdU-stained neuroblasts. A gray scale attribute filter with an opening function of area minimum of 15 pixels and connectivity of 8 was applied to isolate the cell bodies of DsRed-stained neuroblasts, and of an area minimum of 50 pixels and connectivity of 8 to isolate CC3-stained cells. After making a mask of the images, the density of stained cell bodies was assessed by counting the number of cells with the “Analyze Particles” function and normalizing this number to the area of the drawn ROI for the layer. GFAP staining was quantified by the percent area stained within the drawn ROI.

We employed the ImageJ plugin FracLac to assess the morphology of individual binarized microglial and neuroblast cells (Young and Morrison, 2018). 40X confocal images were taken of GFP+ microglia and tdTomato+ neuroblasts in the RMS elbow, and max projections were made of 12 µm (16 slices at 0.75 µm step distance) to collapse the whole cell and its processes. Max projections were obtained for four sections of the RMS elbow per animal, with every cell that had all of its processes contained within the image analyzed. For the GFP channel, a “Despeckle” function was first used to deal with the noise, followed by a “remove outliers” function to target bright outliers with a pixel radius of 2 and threshold of 50. The GFP+ cell was isolated from background using the “default” automatic threshold and binarized. The “close” plugin was then used to connect two pixels if they were separated by up to 2 pixels. To enhance the small features of neuroblasts and remove background in the tdTomato channel, a ‘difference of Gaussians’ was applied to the max projection. The tdTomato+ cell was isolated from background using the “triangle” automatic threshold and binarized before performing fractal analysis.

To perform fractal analysis on an individual cell in the image, a rectangle selection of 100 pixels of both height and width was applied for the ROI of all cells, to ensure that all the cells have the same scale. While fractal shapes are scale-independent, FracLac for ImageJ is dependent on scale, and thus the ROI needs to be consistently sized throughout the data collection (Karperien, 1999; Young and Morrison, 2018). Using the matching max projection photomicrograph as a reference, the paintbrush tool was used to isolate the cell of interest by removing adjacent cell processes and connecting fragmented processes (Young and Morrison, 2018). The isolated cell was then converted to an outline using the “outline” function. Fractal analysis data is gathered via box counting, in which a series of grids of decreasing caliber were systematically laid over an image and the number of boxes containing a pixel counted. The fractal dimension (D_B_) is the relationship between how a pattern’s detail (N) changes with the scale (ε), or resolution, at which the image is considered (Karperien et al., 2013). Fractal analysis was done with the Box Counting in FracLac plugin, and data was collected with 4 box counting orientations (Num G = 4). Lacunarity (Λ) is also calculating using the box counting method and is a measure of cell heterogeneity; cells with low lacunarity are homogenous or rotationally invariant (Karperien et al., 2013). Additional cell shape measurements were generated using the “convex hull” (a polygon that encloses all pixels of the binary cell) and “bounding circle” (the minimum circle that can enclose all pixels.) The span ratio is the ratio of the longest length over the longest length of the convex hull (Morrison et al., 2017). Density is the ratio of the area of cell divided by the area of its convex hull. A full list and explanation of FracLac output data is explained in the FracLac for ImageJ manual (Karperien, 1999). The calculations to obtain the fractal dimension (D_B_), lacunarity (Λ), span ratio and density are best explained in the reference guide provided for FracLac for ImageJ (http://rsb.info.nih.gov/ij/plugins/fraclac/FLHelp/Introduction.htm).

3D rendering to show the accumulation of fluoroliposomes in microglia was performed using IMARIS software using the “Blend” mode (Bitplane, Imaris Viewer, https://imaris.oxinst.com/imaris-viewer).

### Statistical Analysis

All measures across different sections within an animal from the same RMS region were averaged to yield one sample replicate for statistical analysis. Statistical analysis was performed with GraphPad Prism 9.2 for windows (GraphPad Software Inc, San Diego, US, www.graphpad.com). The data was analyzed with a one-way ANOVA followed by Bonferroni’s post-hoc test or a Student t-test as appropriate. *p <0.05; **p < 0.01; ***p<0.001; ****p<0.0001. Data are shown as the mean ± SEM. Specific p values for each experiment are shown in the statistical table (Table 3).

**Table 3.**
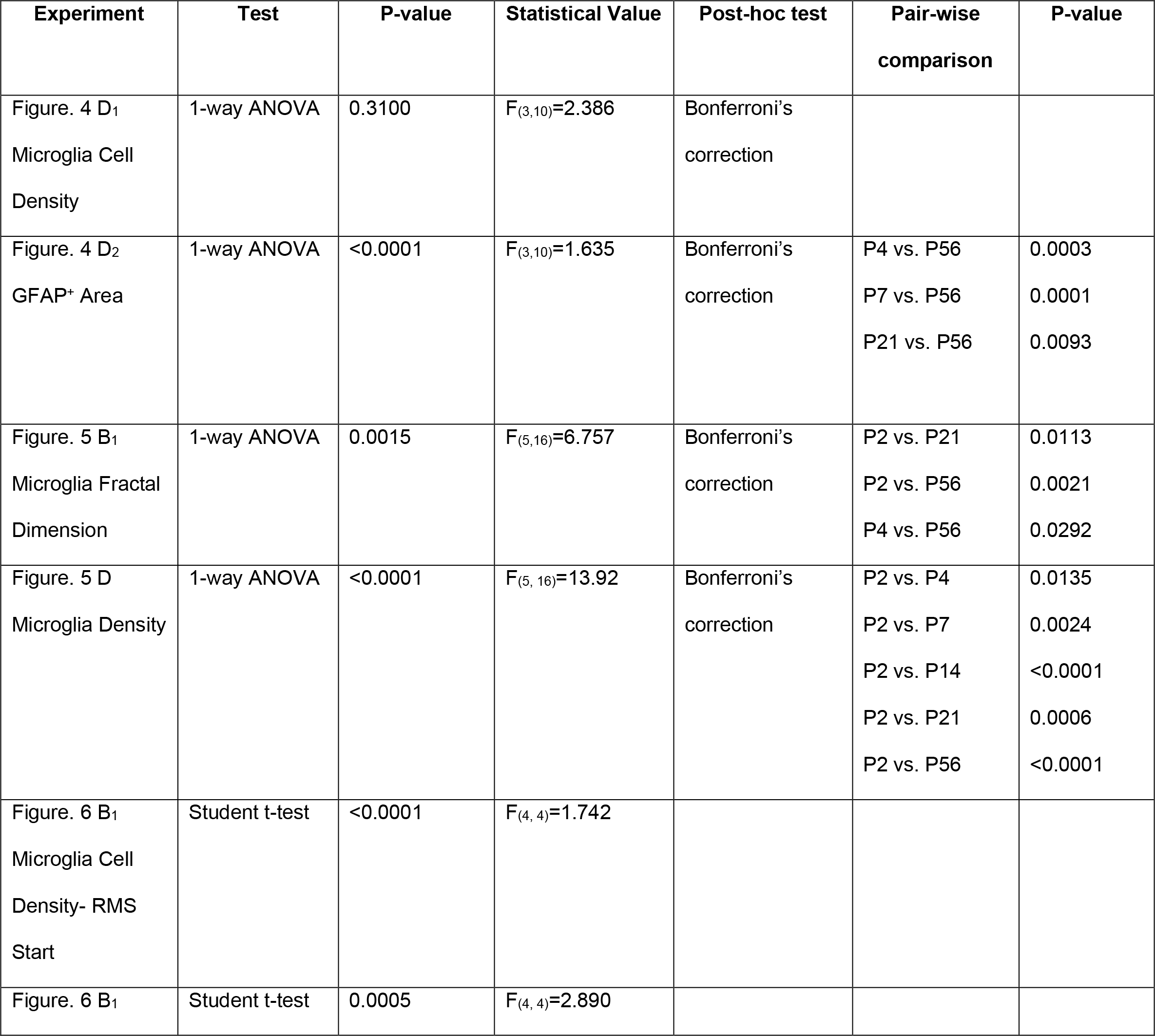
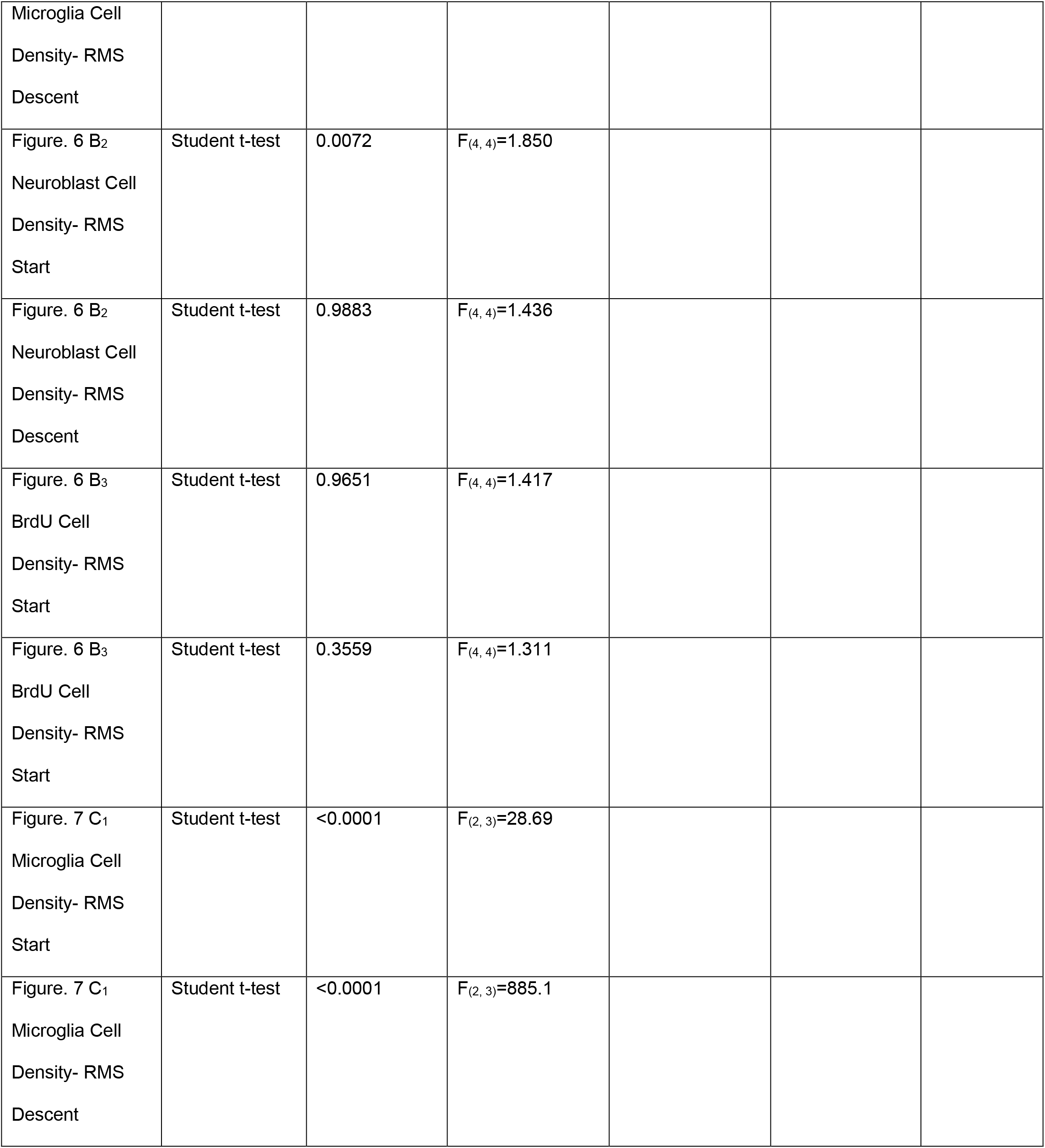
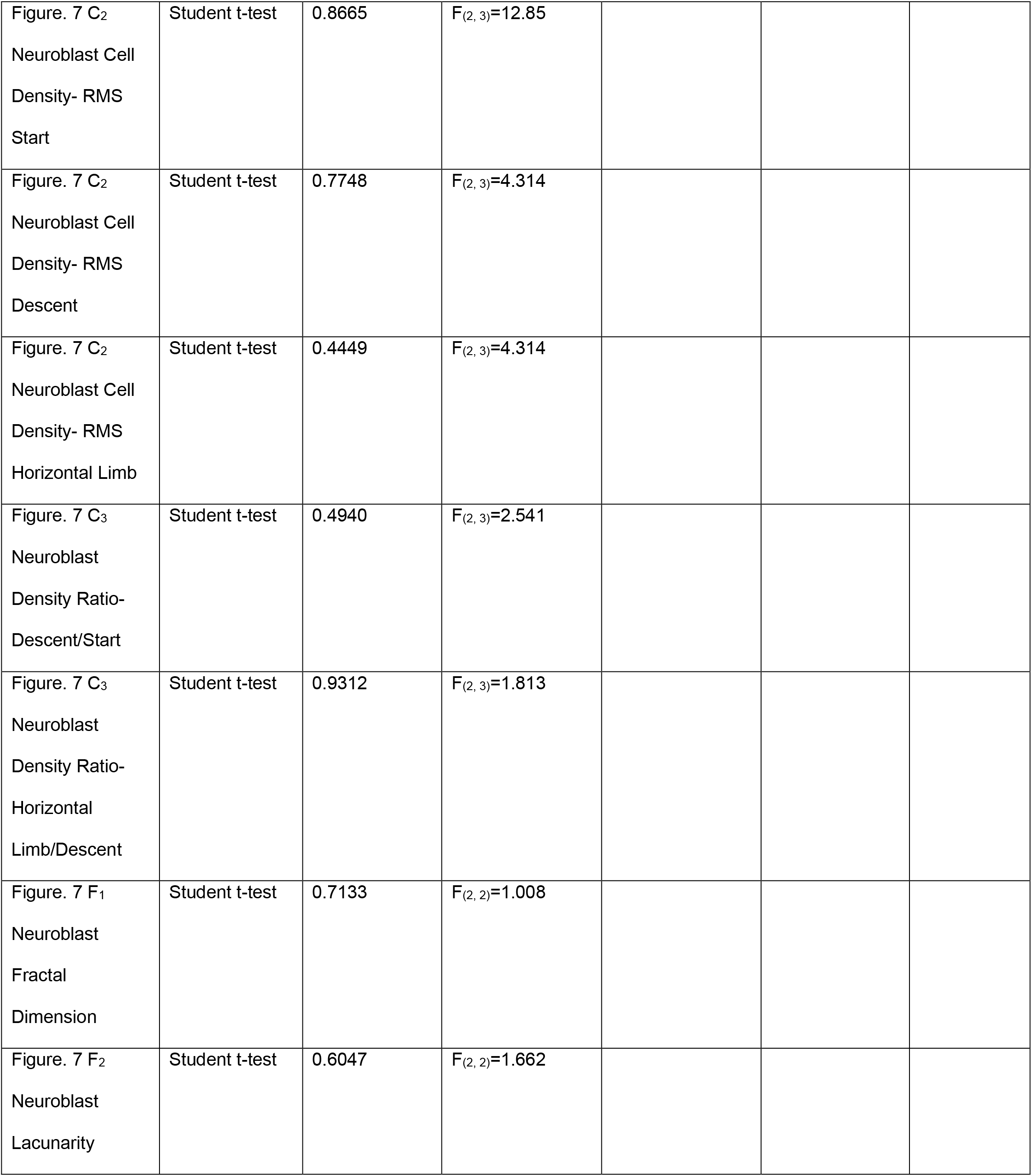
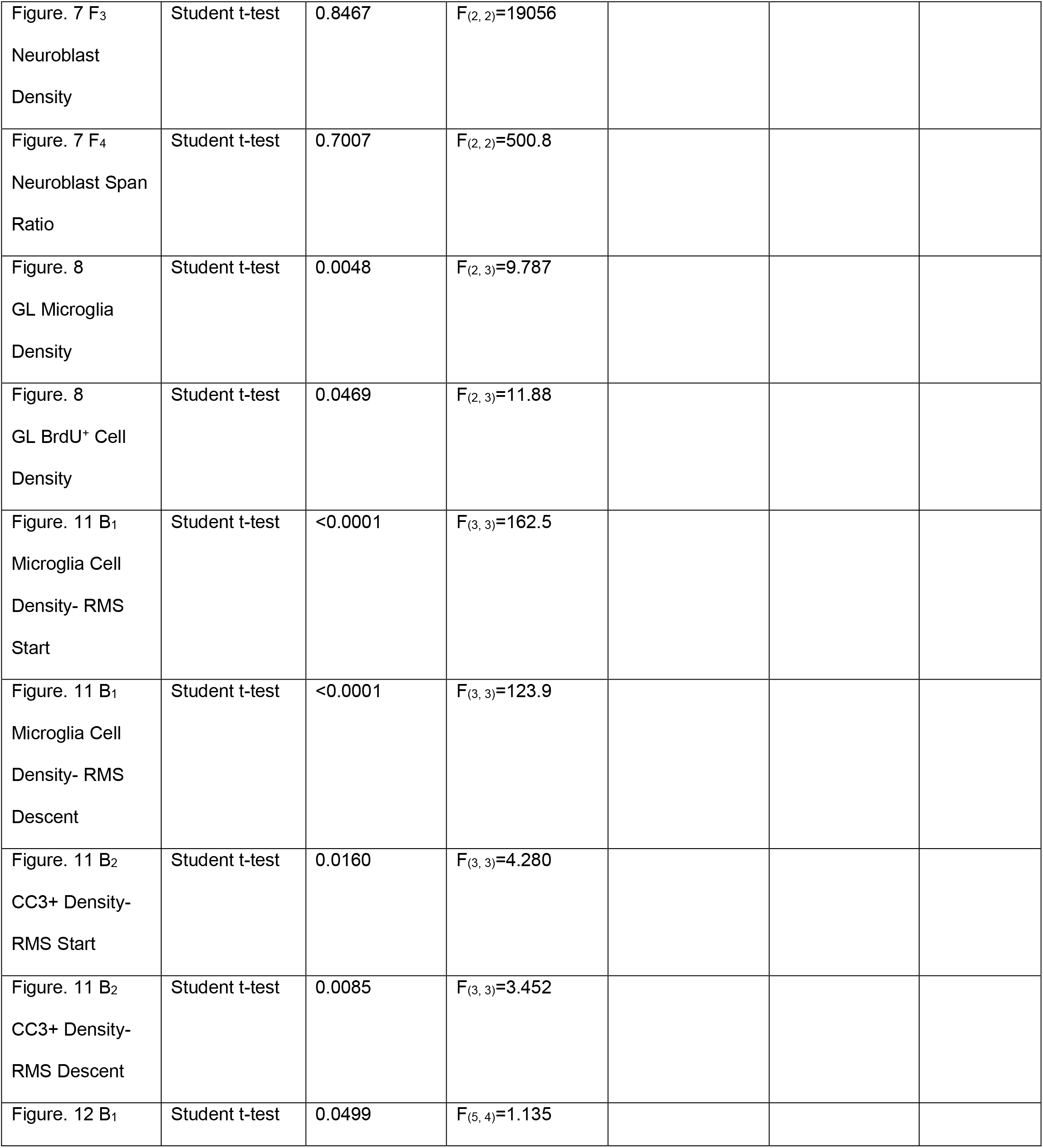
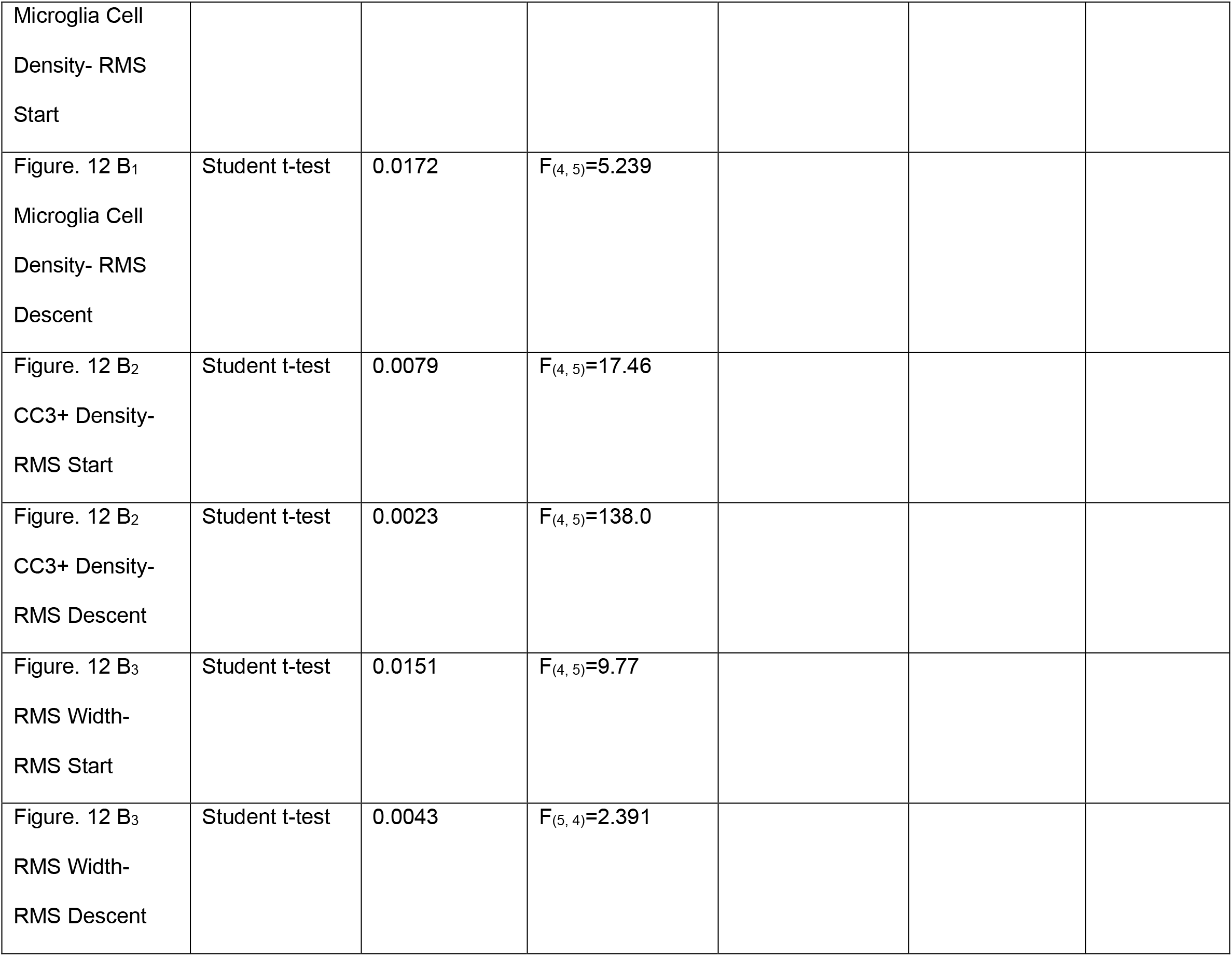
Summary of statistical analysis employed in each experiment. Note - n.s.: not significant.

## RESULTS

### Ameboid microglia are present in the developing RMS prior to the formation of the astrocyte and vascular scaffold

There is active migration of neuroblasts from the SVZ to the OB through the RMS during the first postnatal week, as shown by dense staining of DsRed+ neuroblasts in the RMS of *DCX^DsRed/+^* mice (Fig. 2A). During the first postnatal week microglia line the RMS along its lateral borders (Fig. 2B). Within the RMS, microglia closely appose neuroblasts that were electroporated to express tdTomato (Figure 2B insert). This close interaction between microglia and neuroblasts within the RMS was confirmed with electron microscopy (Figure 2C_1-4_), which showed microglia and their processes (green) intimately associated with migrating neuroblasts. We identified neuroblasts based on the following ultrastructural features: multiple nuclei within the nucleus, a dark cytoplasm with many free ribosomes, and the presence of extracellular spaces between neuroblasts (Doetsch et al., 1997; Matsumoto et al., 2019).

**Figure 2.**
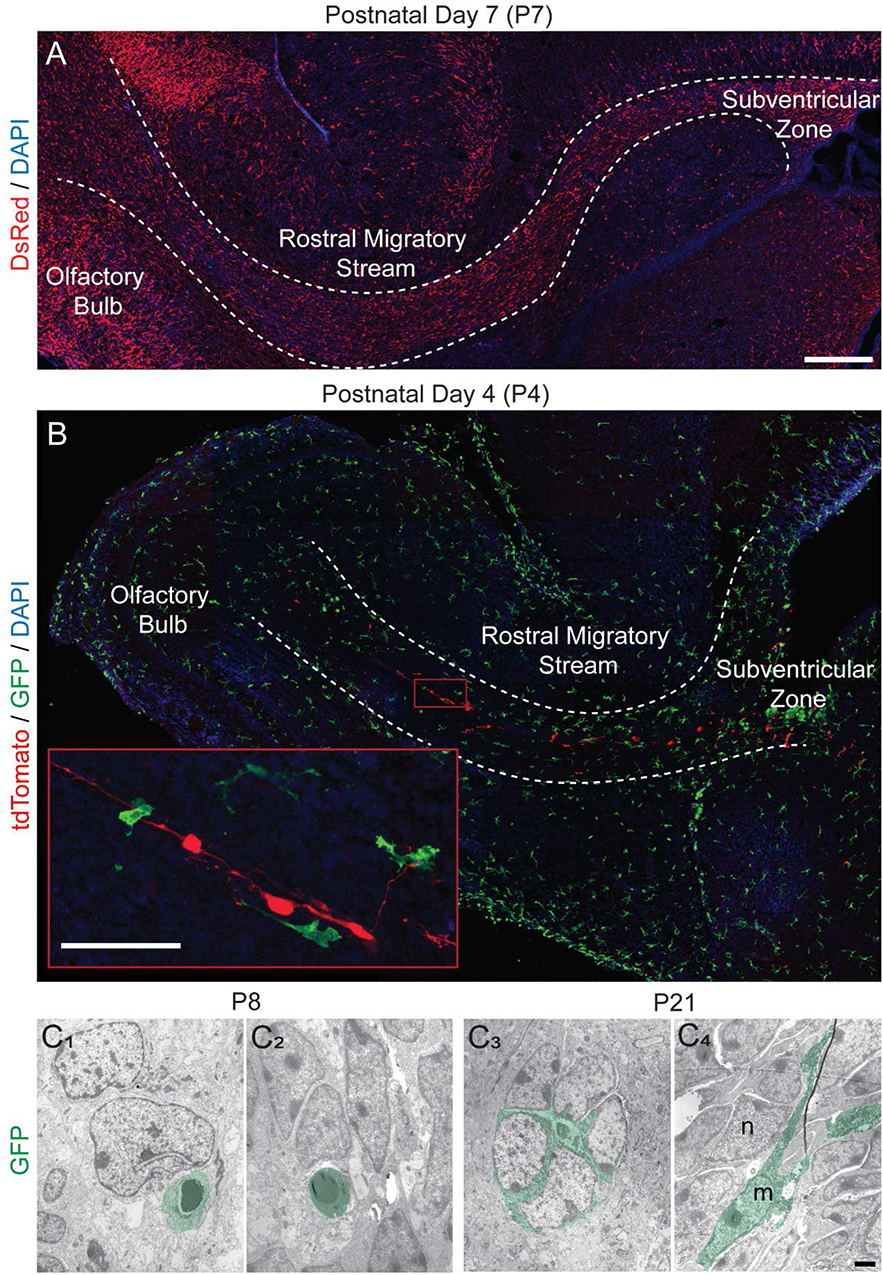
Microglia line the rostral migratory stream (RMS) and contact migrating neuroblasts during early postnatal development. (A) Sagittal section showing neuroblasts (red) in a P7 *DCX^DsRed/+^* mouse. There is active migration of neuroblasts from the subventricular zone (SVZ) to the olfactory bulb (OB) via the RMS in the first postnatal week. Scale bar = 250 μm. (B) Sagittal section showing microglia (green) in the SVZ, RMS and OB in a P4 *CX3CR-1^GFP/+^* mouse. To identify individual migrating neuroblasts (red), the SVZ was electroporated with a tdTomato plasmid at P0 and sacrificed at P4. Microglia line the borders of the RMS and associate with migrating neuroblasts within the core. Scale bar = 50 μm. (C) Ultrathin sections showing GFP-immunopositive microglia soma and processes at the electron microscope level in *CX3CR-1^GFP/+^* mice at P8 and P21. Microglia are pseudo-colored in green. (C_1_-C_2_) Ameboid microglia at P8 contact migrating neuroblasts. (C_3_-C_4_) Microglia processes contact migrating neuroblasts at P21. (C_3_) A microglia between a cluster of migrating neuroblasts. m, microglial cell. n, neuroblast. Scale bar = 2 μm.

Microglia distribute along the RMS prior to the establishment of the vascular and astrocyte scaffold. While CD31+ stained vasculature is observed during the first postnatal week, it does not show the characteristic wrapping of the RMS that forms the scaffold during adulthood (Fig. 3A_1-3_). Numerous tdTomato labeled neuroblasts are seen at a distance from the developing vasculature, suggesting that they migrate independently of the vasculature during this period. An increase in CD31 stained blood vessels was observed at P14, with the beginnings of a network of vasculature forming around the RMS at this age (Fig. 3A_4_). Microglia closely appose endothelial cells in the RMS at P4 and P21 (Fig. 3A_5_, B).

**Figure 3.**
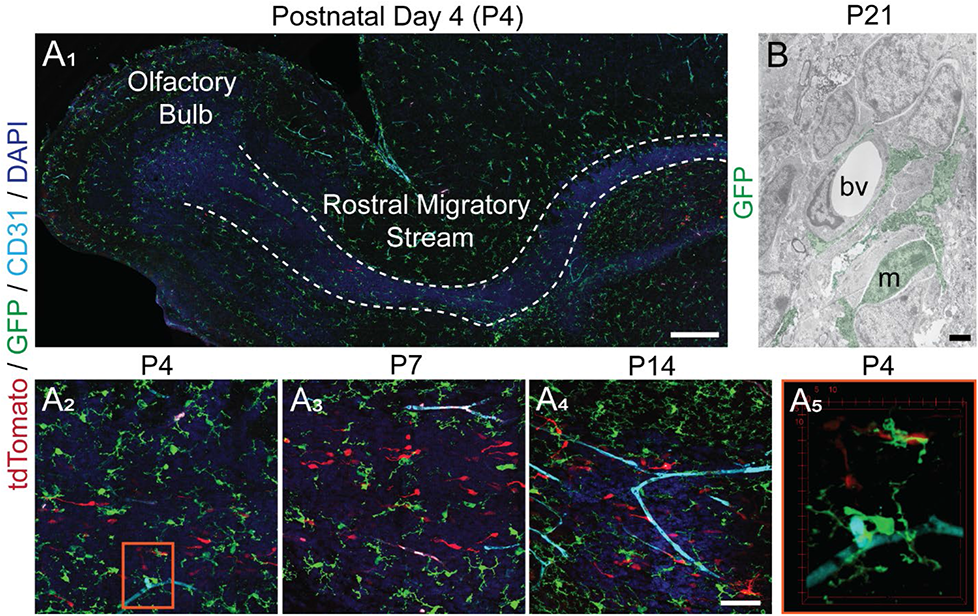
Microglia are present in the rostral migratory stream (RMS) prior to the formation of the vascular scaffold. (A) Sagittal sections of the RMS from *CX3CR-1^GFP/+^* mice electroporated with a tomato plasmid at P0 and sacrificed at ages P4, P7 and P14. Migrating neuroblasts are depicted with tomato, microglia with GFP, and vasculature with CD31. (A_1_) The RMS region was defined by increased density of DAPI nuclear staining, as shown by the white dashed outline. While there is CD31 labeled vasculature in the parenchyma at P4, there is not the stereotypic vascular wrapping of the RMS observed in adulthood. Scale bar = 250 μm. (A_2-4_) Microglia closely associate with migrating neuroblasts in the RMS at P4, P7 and P14. Note that the majority of migrating neuroblasts are not in close vicinity of the developing vasculature during the first postnatal week. Scale bar = 50 μm. (A_5_) The orange outlined inset depicts a volume rendering of microglia interacting with vasculature and the soma of a migrating neuroblast at the RMS elbow at P4. (B) Ultrathin electron micrograph sections showing GFP-immunopositive microglia in a P21 *CX3CR-1^GFP/+^* mouse contacting an endothelial cell of a blood vessel in the RMS. m, microglial cell. bv, blood vessel. Scale bar = 2 μm.

Similarly, an astrocyte tube of thick septate processes that surrounds and infiltrates the RMS is present in the adult mouse (P56) but not during the first postnatal week (P7) (Fig. 4C). Instead, radial glia stained with glial fibrillary acidic protein (GFAP) radiate out from the RMS core during the first postnatal week (Fig. 4A). Microglia organized along the periphery of the RMS during the first postnatal week closely intermingled with the GFAP stained radial glial processes (Fig. 4A). The gradual formation and thickening of astrocyte processes over development in the RMS is reflected in the significant increase in GFAP staining across the elbow between P4 and P56 (p < 0.0001; Fig. 4D_2_). By P56 microglia exhibit a more homogeneous distribution across the RMS (Fig. 4C), reminiscent of the ‘tiling’ observed in the mature cortex (Nimmerjahn et al., 2005). While the microglia distribution changes over the course of postnatal RMS development, there is no significant difference in the density of GFP+ microglia in the RMS between postnatal ages (Fig. 4D_2_).

**Figure 4.**
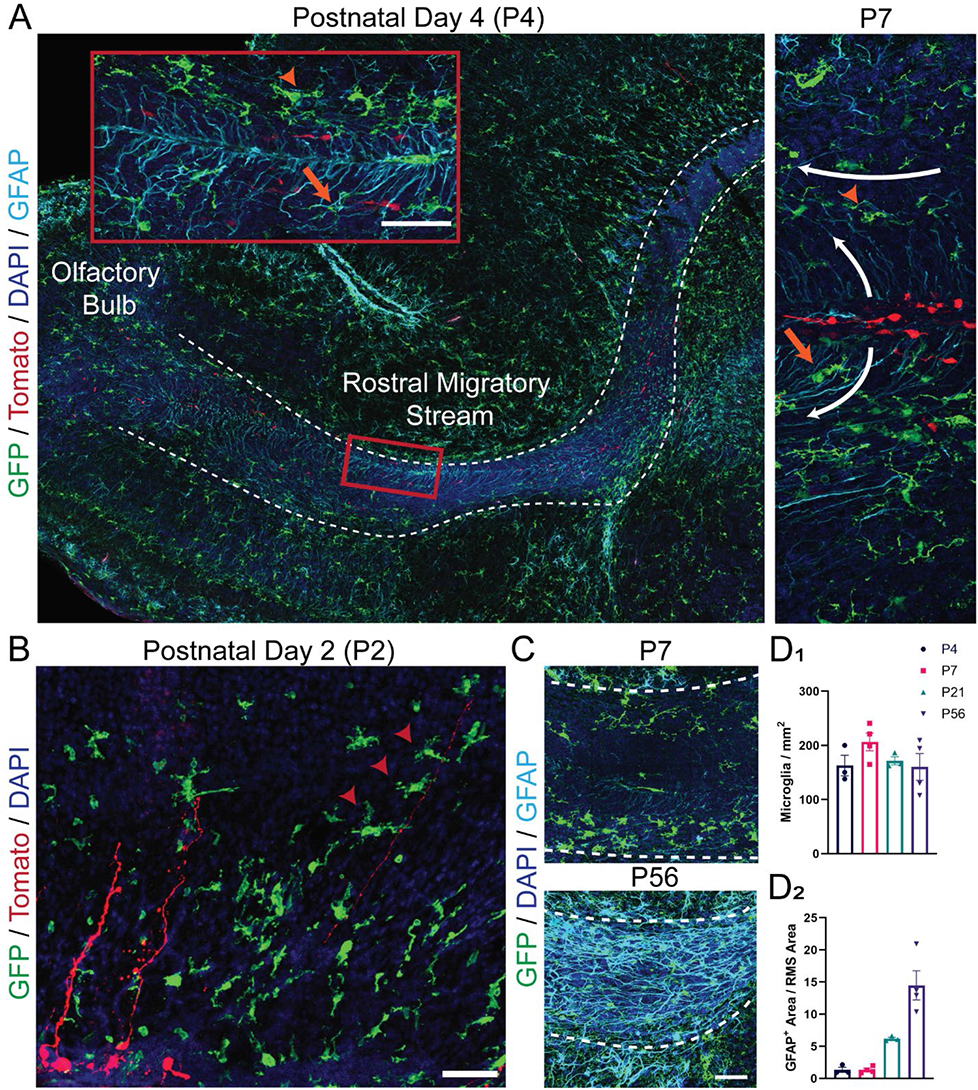
Microglia are present in the rostral migratory stream (RMS) prior to the formation of the astrocytic tube and interact with radial glia. (A) Sagittal sections of the RMS from *CX3CR-1^GFP/+^* mice electroporated with a tomato plasmid at P0 and sacrificed at ages P4 and P7. Migrating neuroblasts are depicted with tomato, microglia with GFP, and radial glia and astrocytes with GFAP. Microglia interact with GFAP stained radial glial processes tangentially oriented at the border of the RMS (orange arrowheads) and with processes radially distributing within its core (orange arrows). Scale bar = 50 μm. (B) Sagittal section of radial glia cell pial fibers extending from the SVZ of a *CX3CR-1^GFP/+^* mouse electroporated with a tomato plasmid at P0 and sacrificed at ages P2. Some microglia interact with pial fibers via a process extension that partially wraps around the fiber in the manner of a “hand-hold” (red arrowheads). Scale bar = 50 μm. (C) The RMS elbow from *CX3CR-1^GFP/+^* mice at P7 and P56. Microglia are represented by GFP (green) and radial glia and astrocytes by GFAP (cyan). Scale bar = 50 μm. Microglia are observed predominately at the periphery of the RMS at P7, where they associate with radial glia processes. While the astrocyte tube is not present at P7, the thick septate processes of GFAP-stained astrocytes are present in the RMS by P56. (D_1_) There is a dramatic increase in GFAP staining between P4 and P56 in the RMS elbow, reflecting the development of the astrocyte tube. (D_2_) Microglia are present in the RMS elbow at similar densities at P4, P7, P21 and P56.

The morphology of microglia in the developing RMS may further give insight into their function. Ameboid microglia are round with short processes and appear to be highly motile and phagocytic (Davalos et al., 2020). Conversely, ramified microglia have long and highly branched processes that continuously sample their local environment (Davalos et al., 2005; Nimmerjahn et al., 2005). Microglia exhibit a greater range of morphologies during the early postnatal period in the RMS, including fewer morphologically complex and more “dense” cells (p < 0.0001; Fig. 5). The “dense” microglia observed at P2 and P4 implicate ameboid morphologies in the RMS during the first postnatal week (Fig. 5D). There is a decrease in very “dense” and ameboid cells by P14, which is accompanied by an increase in cell complexity (Fig. 5). Microglia in the RMS show a gradual increase in cell process complexity between P2 and P56, as demonstrated by the increase in the average fractal dimension (D_B_) of binarized microglia (p < 0.01; Fig. 5B_1_). This increase in the complexity of microglia processes reflects an increased ramification. This change in microglia morphology from ameboid to ramified during RMS development could simply be a manifestation of normal microglial cell development. However, it seems plausible that the functional state of microglia adapt in response to the signals from their local tissue environment (Matcovitch-Natan et al., 2016; Li and Barres, 2018), including cues from the emerging astrocyte and vascular scaffolds.

**Figure 5.**
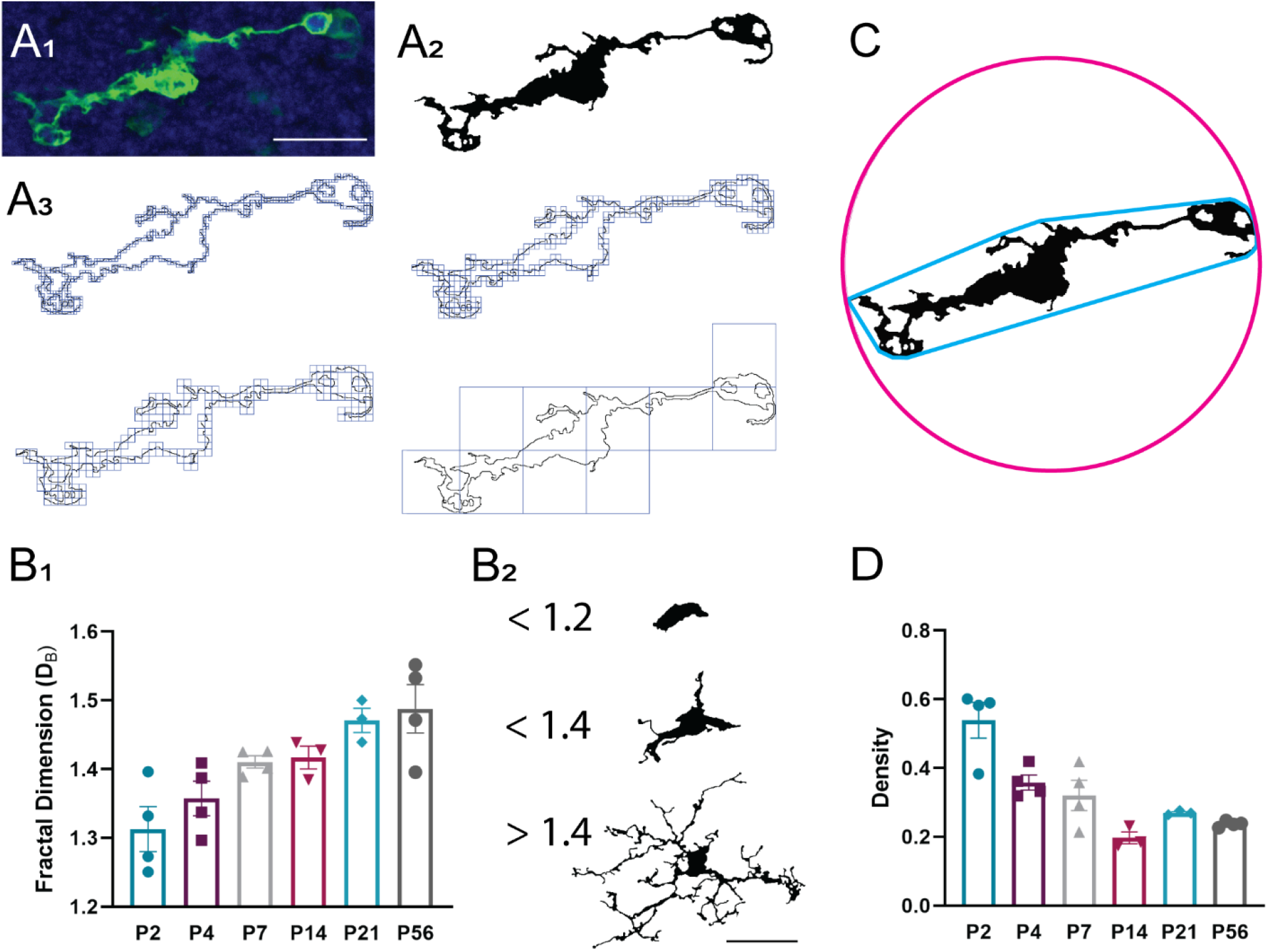
Microglia in the rostral migratory stream exhibit increased average pattern complexity at later postnatal ages. (A_1_) A microglial cell from the RMS of a P4 *CX3CR-1^GFP/+^* mouse. Scale bar = 25 μm. (A_2_) The microglia from (A_1_) segmented and binarized. (A_3_) Fractal analysis data is gathered via box counting, in which a series of grids of decreasing caliber were systematically laid over an image and the number of boxes containing a pixel counted. The fractal dimension (D_B_) is the relationship between how a pattern’s detail (N) changes with the scale (ε) at which the image is considered. (B_1_) There is significantly increased average pattern complexity at later postnatal ages, as shown by increased measures on the fractal dimension. (B_2_) Example microglia shapes and their corresponding fractal dimension D_B._ Scale bar = 25 μm. (C) Microglia cell “density” is measured by dividing the area of the cell by the area of its convex hull. The convex hull is in cyan, which is created by connecting a series of straight segments that enclose all the pixels of the binary image. The bounding circle (depicted in magenta) is the smallest circle enclosing all the pixels. (D) There is a decrease in the proportion of “dense” and more “ameboid” cells at later postnatal ages. 8 – 20 microglia per animal.

### Neuroblast migratory capacity is independent of microglia in the developing rostral migratory stream

The presence of microglia in the RMS during the first postnatal week prompted the hypothesis that microglia may facilitate neuroblast migration in the absence of the vascular and astrocyte scaffold. We first investigated whether microglia depletion affects the density of neuroblasts migrating down the RMS; neuroblast clumping within the RMS following microglia depletion would implicate a lack of forward migration. Microglia depletion was carried out in *CX3CR-1^GFP/+^;DCX^DsRed/+^* transgenic mice to allow identification of microglia (GFP) and neuroblasts (DsRed) (Fig. 6A_1_). To further analyze the density of migrating neuroblasts, 25 mg/kg of BrdU was injected at P0 to label actively diving neuroblasts (Fig. 6A_2_). Microglia depletion was initiated two hours later with an injection of clodronate-filled liposomes in the cerebral lateral ventricles. PBS-filled liposomes injected in littermates served as a control. Mice were killed at P3 to determine the density of BrdU- and DsRed-labeled neuroblasts in the RMS start and descent following microglia depletion. There was a significant reduction in the density of microglia in both the RMS start and RMS descent in mice injected with liposomal clodronate as compared to littermates injected with liposomal PBS (p < 0.001; Fig. 6B_1_). However, microglia depletion was not accompanied by a difference in the density of either DsRed- or BrdU-labeled neuroblasts in the RMS descent (p = n.s.; Fig. 6B_2_-B_3_). While there was a significant increase in the density of DsRed-labeled cells in the RMS start following liposomal clodronate treatment (p < 0.01; Fig. 6B_2_), there was no significant difference in the density of BrdU-labeled neuroblasts in the RMS start (Fig. 6B_3_). The absence of difference in BrdU labeling suggests that neurogenesis was not significantly altered following transient microglia depletion. The increase in DsRed staining may reflect a decrease in microglia phagocytosis of neural stem cells in the SVZ, which was not captured by the temporally sparse BrdU labeling of newly born neuroblasts. The similar density of neuroblasts migrating down the RMS descent following liposomal clodronate treatment indicates that neuroblasts are not stalled in their migration toward the OB in the absence of microglia.

**Figure 6.**
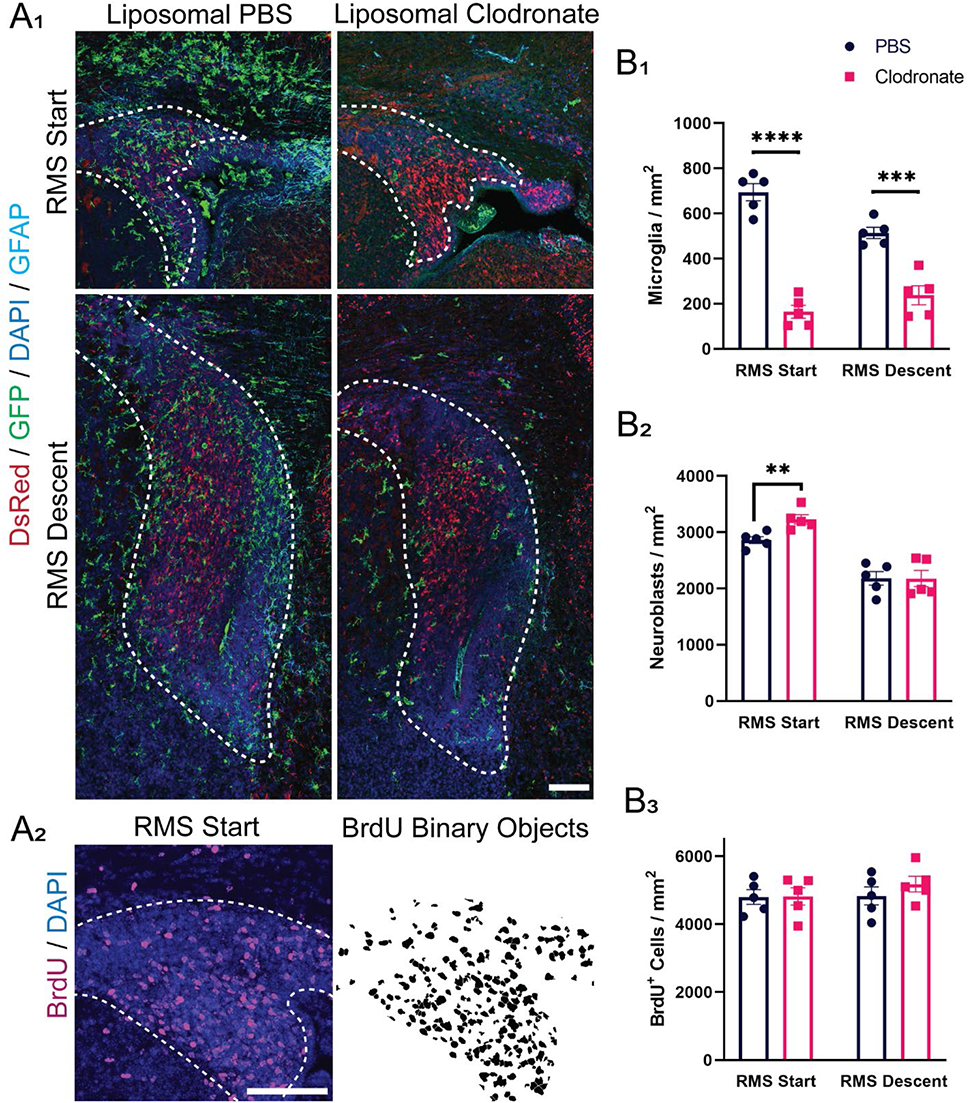
Microglia depletion after liposomal clodronate injection does not impact neuroblast density in the rostral migratory stream (RMS). (A_1_) Coronal sections of the RMS start and descent from P3 *CX3CR-1^GFP/+^;DCX^DsRed/+^* littermates following injection with liposomal PBS at P0 and subcutaneous 25 mg/kg BrdU injection two hours later. The RMS region was defined by increased density of DAPI nuclear staining, as shown by the white dashed outline. Scale bar = 100 μm. (A_2_) GFP-, DsRed-, and BrdU-stained cells were segmented and binarized with Fiji software. Scale bar = 100 μm. (B_1_) Three days after injection of liposomal clodronate, there was a significant decrease in microglia density in both the RMS start and RMS descent. (B_2_) Microglia depletion was not accompanied by a corresponding decrease in the density of migrating neuroblasts in the RMS descent, as shown by the density of DsRed-labeled cells in this region. There was a significant increase in the density of DsRed-labeled cells in the RMS start following liposomal clodronate treatment, potentially reflecting a decrease in microglia phagocytosis of neural stem cells in the subventricular zone. (B_3_) There was no significant difference in the density of newly generated BrdU-labeled neuroblasts in either the RMS start or descent between liposomal PBS and liposomal clodronate treated littermates. **p < 0.01; ***p<0.001; ****p<0.0001.

To further examine whether microglia promote neuroblast migration we asked if microglia depletion impacts the distance migrated by individual neuroblasts in the RMS. To accomplish this we used an alternative model of inducible microglia depletion. Homozygous *Cx3cr1^CreER/CreER^* were crossed with heterozygous *ROSA26^eGFP-DTA/+^* mice to create *Cx3cr1^CreER/+^; ROSA26^eGFP-DTA/+^* and *Cx3cr1^CreER/+^; ROSA26^+/+^* littermates. Following tamoxifen injection, the toxic DTA subunit is expressed in *Cx3cr1^CreER/+^; ROSA26^eGFP-DTA/+^* mice, leading to the ablation of microglia (Fig. 1). To label individual neuroblasts and evaluate the distance traveled, we electroporated a tdTomato plasmid into the subventricular zone at P0. We injected tamoxifen i.p. two hours later to selectively deplete microglia in those littermates that express the toxic DTA subunit. Littermates that did not express the DTA subunit served as controls. We killed the mice at P3 and assessed the density of tdTomato-labeled neuroblasts in the RMS start, descent and horizontal limb (Fig. 7A-C) and their morphology (Fig. 7D-E) following microglia depletion.

**Figure 7.**
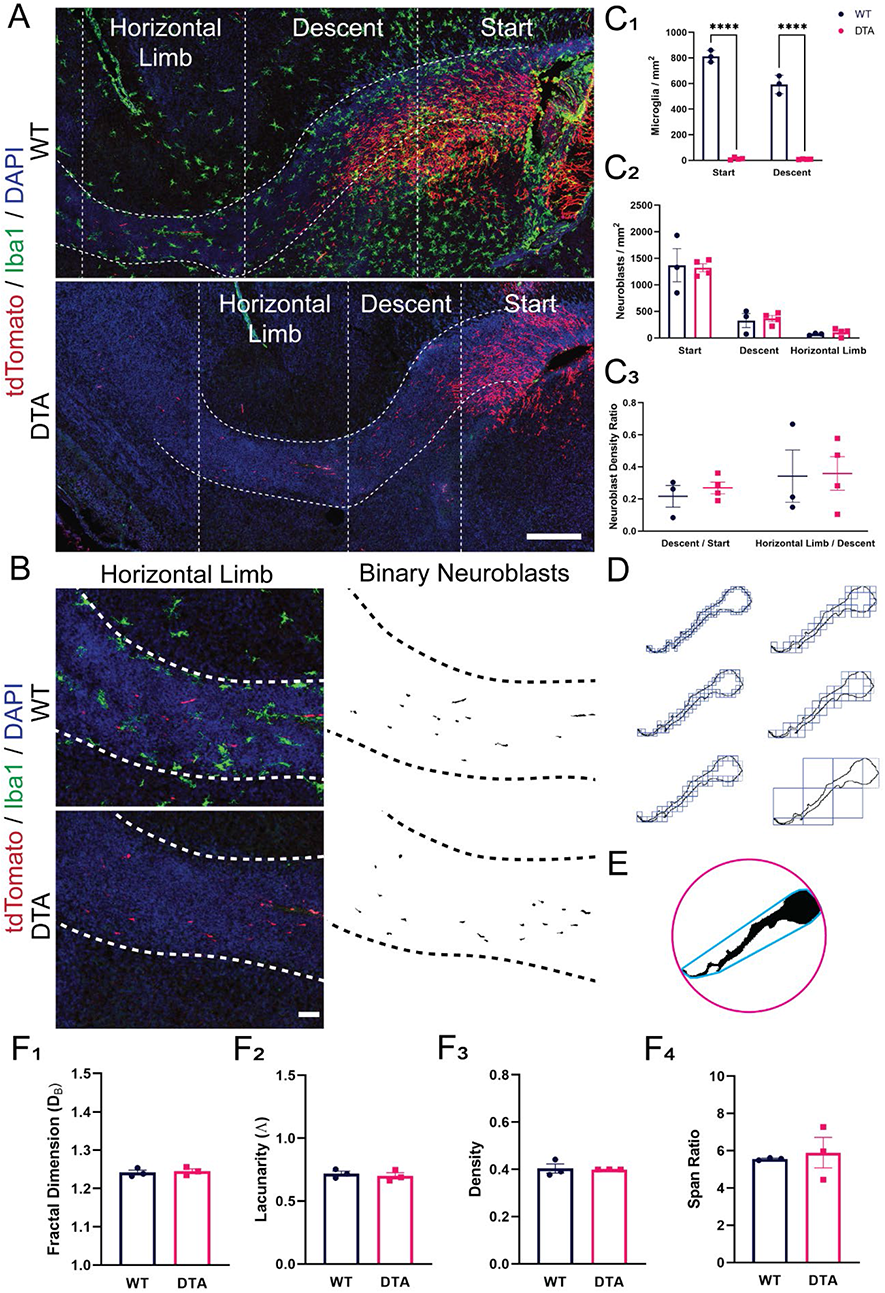
Microglia depletion does not impact the migratory capacity of neuroblasts in the rostral migratory stream (RMS). (A) Sagittal sections of the RMS following i.p. tamoxifen injection in *Cx3cr1^CreER/+^; ROSA26^+/+^* (top) and in *Cx3cr1^CreER/+^; ROSA26^eGFP-DTA/+^* (bottom) littermates at P0 and sacrifice at P3. Migrating neuroblasts (red) were labeled by electroporation of a tdTomato plasmid in the subventricular zone (SVZ) at P0. Microglia line the rostral migratory stream (RMS) and contact migrating neuroblasts within it at P3 (top). Scale bar = 250 μm. (B) tdTomato-labeled neuroblasts were segmented and binarized with Fiji software. The RMS region was defined by increased density of DAPI nuclear staining, as shown by the white dashed outline. Scale bar = 100 μm. (C) There was a significant decrease in the density of Iba1-labeled microglia in the RMS following tamoxifen injection in DTA mice as compared to WT littermates. However, this was not accompanied by a difference in the density of tdTomato-labeled neuroblasts in the RMS start, descent, or horizontal limb. There was also no difference in the ratio of neuroblasts that reached the RMS descent / RMS start, or the ratio of neuroblasts that reached the horizontal limb / the number that were still traveling the descent. (D) A segmented and binarized neuroblast. Fractal analysis data is gathered via box counting, in which a series of grids of decreasing caliber were systematically laid over an image and the number of boxes containing a pixel counted. (E) A binarized neuroblast showing the convex hull used for morphology analysis. The convex hull is in cyan, which is created by connecting a series of straight segments that enclose all the pixels of the binary image. The bounding circle (depicted in magenta) is the smallest circle enclosing all the pixels. (F_1_) There was no significant difference in neuroblast pattern complexity following microglia depletion, as shown by similar measures on the fractal dimension (D_B_). (F_2_) There was no significant difference in the heterogeneity of binarized neuroblast cells following microglia depletion, as shown by measures of lacunarity (Λ). (F_3_) There was no difference in the “density” of neuroblast morphology following microglia depletion (WT vs DTA p = 0.8467; Student t-test). Density is measured by dividing the area of a cell by the area of its convex hull (F_4_) There was no difference in the “span ratio” of neuroblasts in the RMS following microglia ablation. The span ratio is the ratio of the major over minor axes of the convex hull. 9 – 13 neuroblasts per animal. ****p<0.0001.

Tamoxifen injection at P0 significantly depleted microglia three days later in both the RMS start and RMS descent (p < 0.0001; Fig. 7C_1_). However, this was not accompanied by a difference in the density of neuroblasts in the RMS start, descent, or horizontal limb following microglia depletion in *Cx3cr1^CreER/+^; ROSA26^eGFP-DTA/+^* as compared to *Cx3cr1^CreER/+^; ROSA26^+/+^* littermates (p = n.s.; Fig. 7C_2_). There was also no difference in the ratio of tdTomato-labeled neuroblasts that reached the RMS descent over those in the RMS start, or neuroblasts that reached the horizontal limb compared to those still in the RMS descent for each animal (p = n.s.; Fig. 7C_3_). The successful migration of neuroblasts to the more distal segments of the RMS following microglia depletion suggests that microglia are not necessary for neuroblast migration in the RMS. We next asked if microglia depletion influenced neuroblast morphology, potentially hinting at more subtle disorganization of neuroblast migratory patterns (Fig. 7D-E). However, there was no significant difference in neuroblast pattern complexity (p = n.s.; Fig. 7F_1_), heterogeneity (p = n.s.; Fig. 7F_2_), density (p = n.s.; Fig. 7F_3_) or span ratio (p = n.s.; Fig. 7F_4_) following microglia ablation. The similar morphologies of neuroblasts traversing the RMS further indicates that microglia ablation does not impact the migratory capacity of neuroblasts.

### Microglia depletion may affect neuroblast radial migration in the olfactory bulb

Microglia may promote neuroblast detachment from the RMS and radial migration in the OB, as opposed to modulating the tangential migration of neuroblasts in the RMS. To analyze how microglia may impact the radial migration of neuroblasts to the OB glomerular layer (GL), *CX3CR-1^GFP/+^* mice were injected with clodronate-filled liposomes in the cerebral lateral ventricles at P1. PBS-filled liposomes injected in littermates served as a control. 50 mg/kg of BrdU was subsequently injected at P1 to label actively diving neuroblasts (Fig. 8A). Mice were killed at P4 to determine the density of BrdU labeled neuroblasts in the GL following microglia depletion (Fig. 8B).

**Figure 8.**
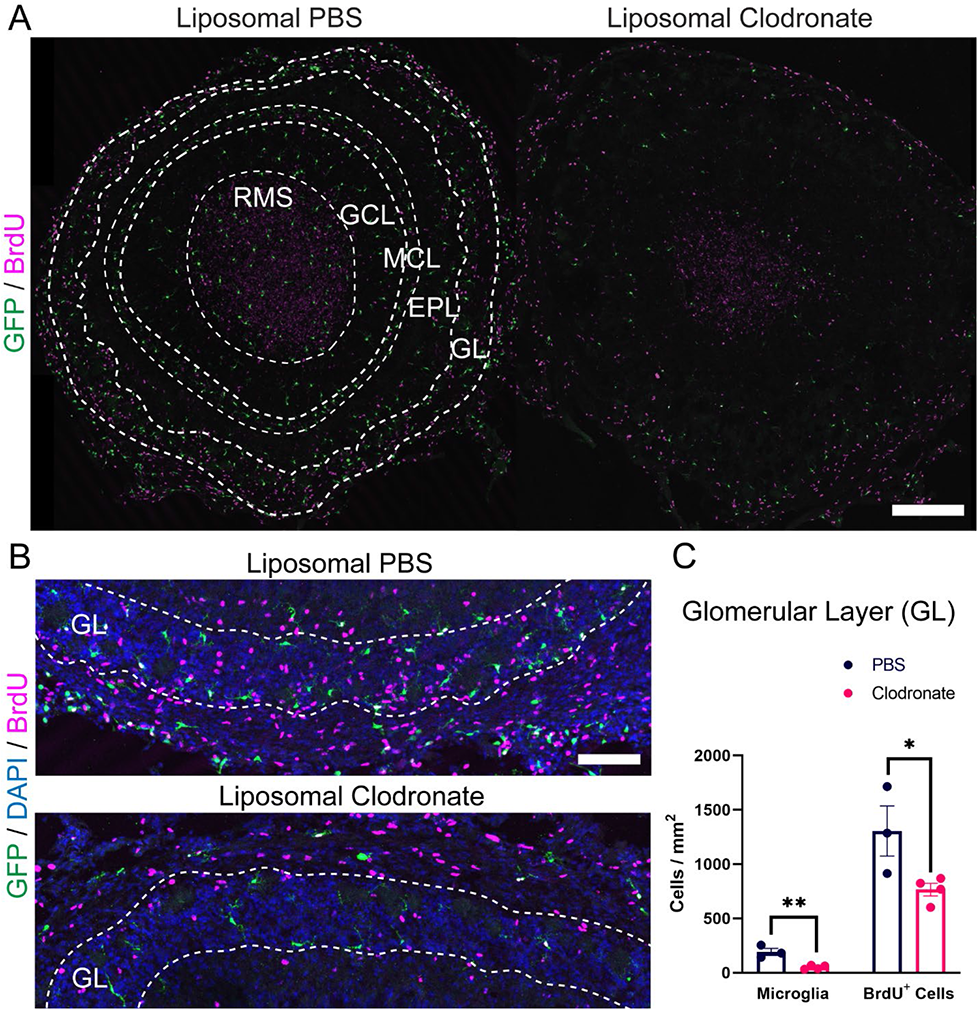
Microglia depletion in the olfactory bulb (OB) after injection with liposomal clodronate decreases BrdU density in the glomerular layer (GL). (a) Coronal sections of whole OBs in *CX3CR-1^GFP/+^* littermates following injection with PBS liposomes (left) and clodronate liposomes (right). Note the laminar distribution of microglia in the mitral cell layer. RMS, rostral migratory stream. GCL, granule cell layer. MCL, mitral cell layer. EPL, external plexiform layer. GL, glomerular layer. Scale bar = 250 μm. (b) The GL region was determined by DAPI nuclear staining, as shown by the dashed white outlines of the GL. Scale bar = 100 μm. (c) There was a significant decrease in the density of GFP labeled microglia in the GL following injection with liposomal clodronate, which was accompanied by a decrease in the density of BrdU labeled cells in the GL. *p <0.05; **p < 0.01.

Liposomal clodronate treatment at P1 was associated with a significant decrease in microglia in the GL at P4 (p < 0.01; Fig. 8C). This was accompanied by a significant decrease in the density of BrdU-labeled cells in the GL (p < 0.05; Fig. 8C). These findings suggest that microglia may promote neuroblast detachment from the RMS, neuroblast radial migration in the GL, or neuroblast survival and differentiation once they reach their final destination in the OB.

### Microglia phagocytosis in the developing rostral migratory stream

The ameboid morphology of microglia in the early postnatal RMS led us to consider a role for microglia phagocytosis in regulating neuroblast number or eliminating aberrantly migrating neuroblasts. While there is little evidence of neuroblast phagocytosis in the adult RMS (Ribeiro Xavier et al., 2015), recent data suggests that specialized microglia exist during the early postnatal period that regulate neurodevelopment through phagocytosis. Microglia express CLEC7A along the RMS in the early postnatal period (Li et al., 2019), while microglial expression of receptor tyrosine kinase MERTK is upregulated on the surface of embryonic microglia compared with the adult, suggesting that receptor-mediated surveillance for phagocytosis may be important in the developing brain (Rosin et al., 2021).

We capitalized on these recent findings to ask if microglia in the RMS exhibit the phagocytic markers CLEC7A, MERTK and CD68 during early postnatal development (Fig. 9A). Microglia expressed phagocytic markers while “hugging” neuroblasts in the corpus callosum of P7 *CX3CR-1^GFP/+^;DCX^DsRed/+^* mice (Fig. 9B), consistent with the phagocytic capacity observed in microglia in this region in previous studies (Li et al., 2019). Clusters of microglia expressing the phagocytic markers CD68, CLEC7A and MERTK were also found wrapping DsRed+ neuroblasts in the RMS start and ventral border of the RMS elbow (Figure 9B-D). Occasional microglia expressing phagocytic markers were also found to wrap migrating neuroblasts in the core of the RMS descent and horizontal limb (Fig. 9B).

**Figure 9.**
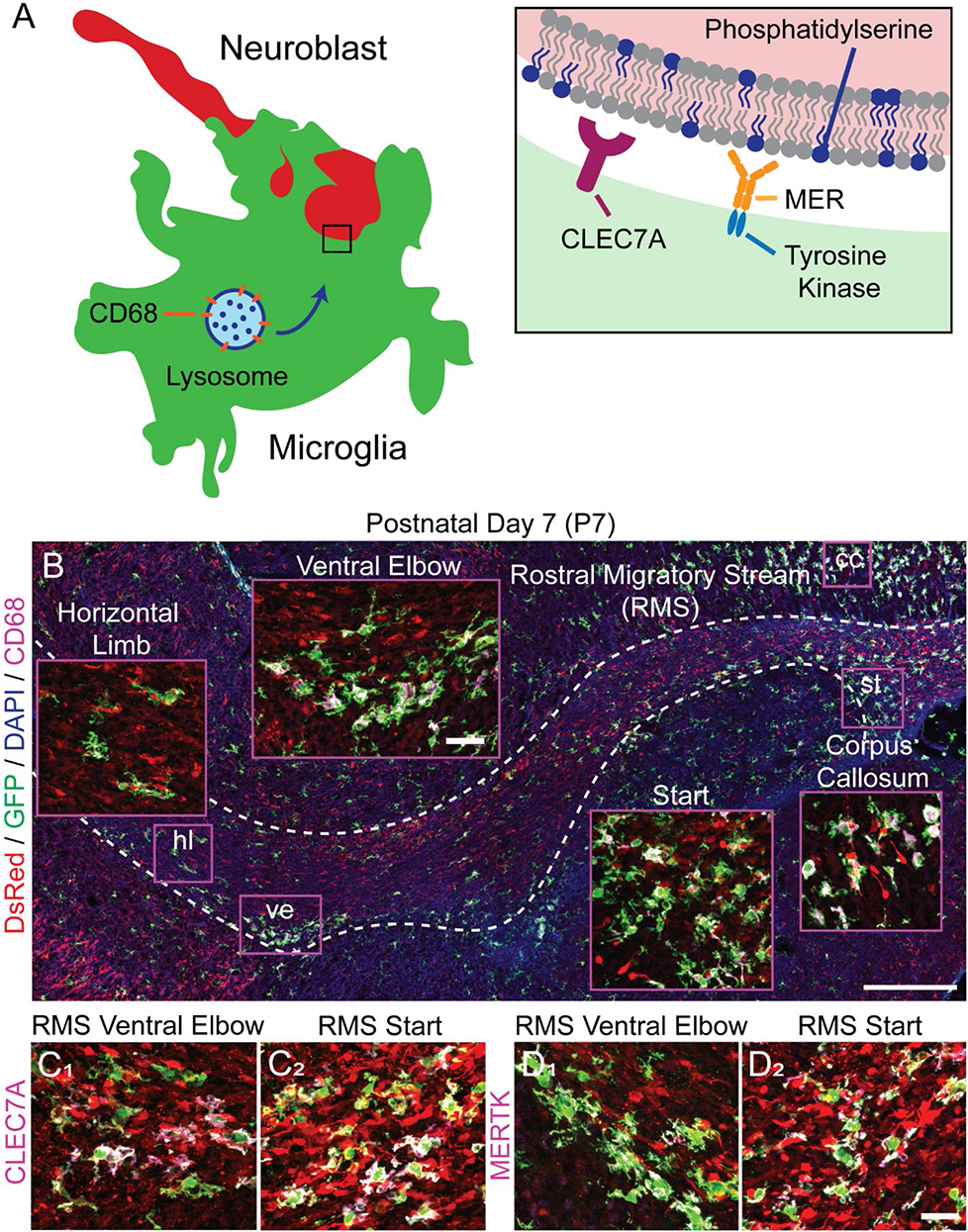
Microglia phagocytose neuroblasts in the postnatal rostral migratory stream (RMS). (A) Illustration of microglia phagocytic markers. CD68 is a lysosome-associated transmembrane glycoprotein that is upregulated with phagocytosis. CLEC7A, or Dectin-1, is a transmembrane pattern-recognition receptor that triggers phagocytosis and the release of reactive oxygen species (ROS). MERTK is a member of the TAM (Tyro3, Axl, and Mertk) family of receptor tyrosine kinases that recognize soluble bridging proteins that bind externalized phosphatidylserine, including GAS6 and protein S. (B) Sagittal section showing the interactions between microglia (green) and neuroblasts (red) in the RMS (dashed white outline) of a P7 *CX3CR-1^GFP/+^;DCX^DsRed/+^* mouse. Microglia exhibited the phagocytic marker CD68 while wrapping neuroblasts in the RMS, indicating active phagocytosis of neuroblasts. Microglia CD68 expression was marked at the start of the RMS and at the ventral border of the elbow. CD68+ microglia also wrapped neuroblast radially migrating from the subventricular zone through the corpus callosum and start of the RMS. cc, corpus callosum. st, start. ve, ventral elbow. Scale bar = 250 μm. Scale bar of magnified inset = 25 μm. Microglia also expressed the phagocytic markers CLEC7A (C) and MERTK (D) while wrapping neuroblasts at the start of the RMS (C_2_, D_2_) and at the ventral border of the RMS elbow (C_1_, D_1_). Scale bar = 25 μm.

We next sought to elucidate whether microglia monitor and phagocytose antigens in the RMS. To demonstrate the phagocytic capacity of microglia throughout the RMS, fluorescent liposomes containing the fluorescent dye Dil (Fluoroliposome) were injected into the cerebral lateral ventricles of P1 CX3CR-1^GFP/+^ mice and killed at P4. Fluoroliposomes accumulated within microglia to a high degree in the SVZ but were also observed in distal regions of the RMS (Fig. 10). Microglia thus appear to monitor and phagocytose extracellular material within the early postnatal RMS.

**Figure 10.**
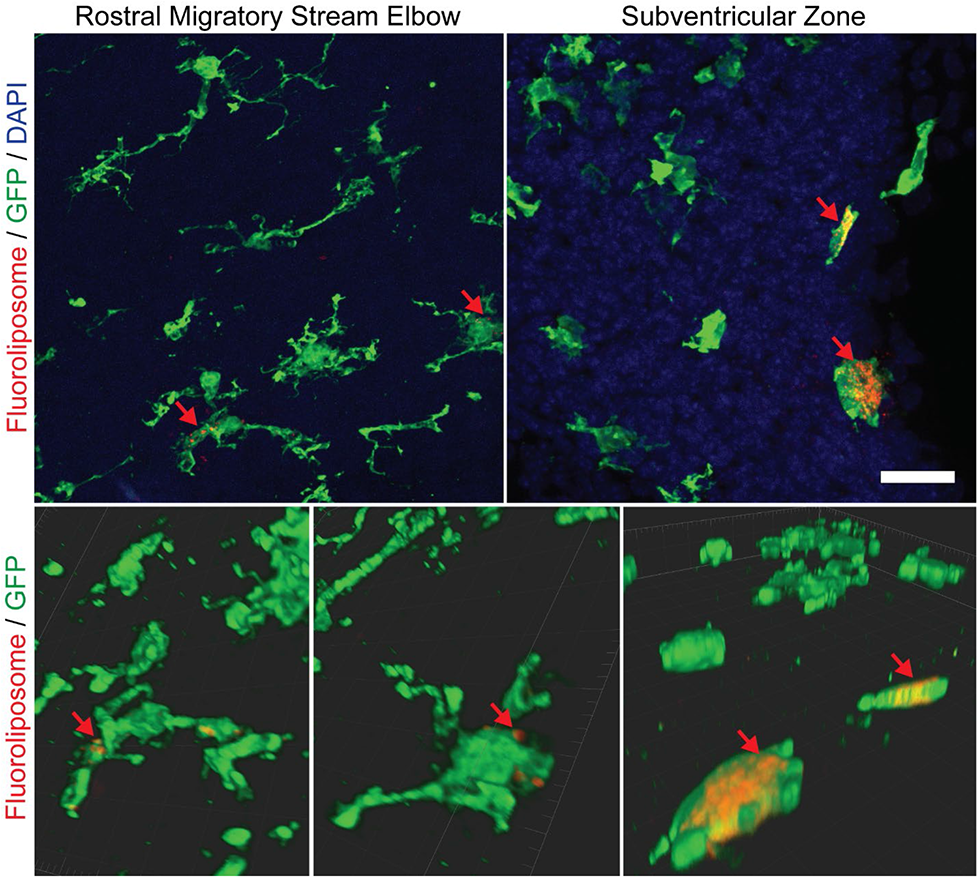
Fluoroliposomes accumulate within microglia in the subventricular zone (SVZ) and rostral migratory stream (RMS). The panels in the top row show sagittal sections of the SVZ and RMS elbow in *CX3CR-1^GFP/+^* mice at P4 after injection of flouroliposome-Dil into the lateral ventricles at P1. Fluoroliposome-Dil have excitation/emission spectra of 550 / 565. The bottom row depicts 3D volume rendering of select microglia with Imaris software. Red arrows indicate flouroliposomes accumulating within GFP+ microglia in the SVZ (right) and ventral border of the RMS (left). Scale bar = 25 μm.

To functionally demonstrate a role for microglia phagocytosis in the developing RMS, we ablated microglia for 3 and 14 days postnatally to evaluate if there was an accumulation of neuroblasts and apoptotic cells in the RMS. *Cx3cr1^CreER/+^; ROSA26^eGFP-DTA/+^* and *Cx3cr1^CreER/+^; ROSA26^+/+^* control littermates were injected with TMX i.p. starting at P0 and killed at P3 or injected with TMX every three days subsequently until sacrifice at P14. To label microglia, neuroblasts and apoptotic cells, immunohistochemistry was performed with primary antibodies against Iba1 (not shown), doublecortin (DCX) and cleaved caspase-3 (CC3) (Fig. 11A, 12A). There was a significant decrease in microglia density in *Cx3cr1^CreER/+^; ROSA26^eGFP-DTA/+^* animals compared to *Cx3cr1^CreER/+^; ROSA26^+/+^* control littermates after both 3 days (RMS start p < 0.0001; RMS descent p < 0.0001; Fig. 11B_1_) and 14 days of TMX treatment (RMS start p < 0.05; RMS descent p < 0.01; Fig. 12 B_1_). Microglia depletion was associated with a significant increase in the accumulation of CC3+ apoptotic cells in the RMS start and RMS descent at both P3 (RMS start p < 0.05; RMS descent p < 0.001; Fig. 11B_2_) and P14 (RMS start p < 0.01; RMS descent p < 0.01; Fig. 12B_2_). Interestingly, an extended period of microglia depletion was also associated with an increase in RMS width at both the RMS start and descent (RMS start p < 0.05; RMS descent p < 0.01; Fig. 12B_3_). The expanded RMS width may reflect an increase in the number of neuroblasts within the RMS following microglia depletion. The accumulation of apoptotic neuroblasts and wide RMS following microglia depletion suggests that microglia may restrict the number of migrating neuroblasts or selectively eliminate those showing aberrant migration within the RMS.

**Figure 11.**
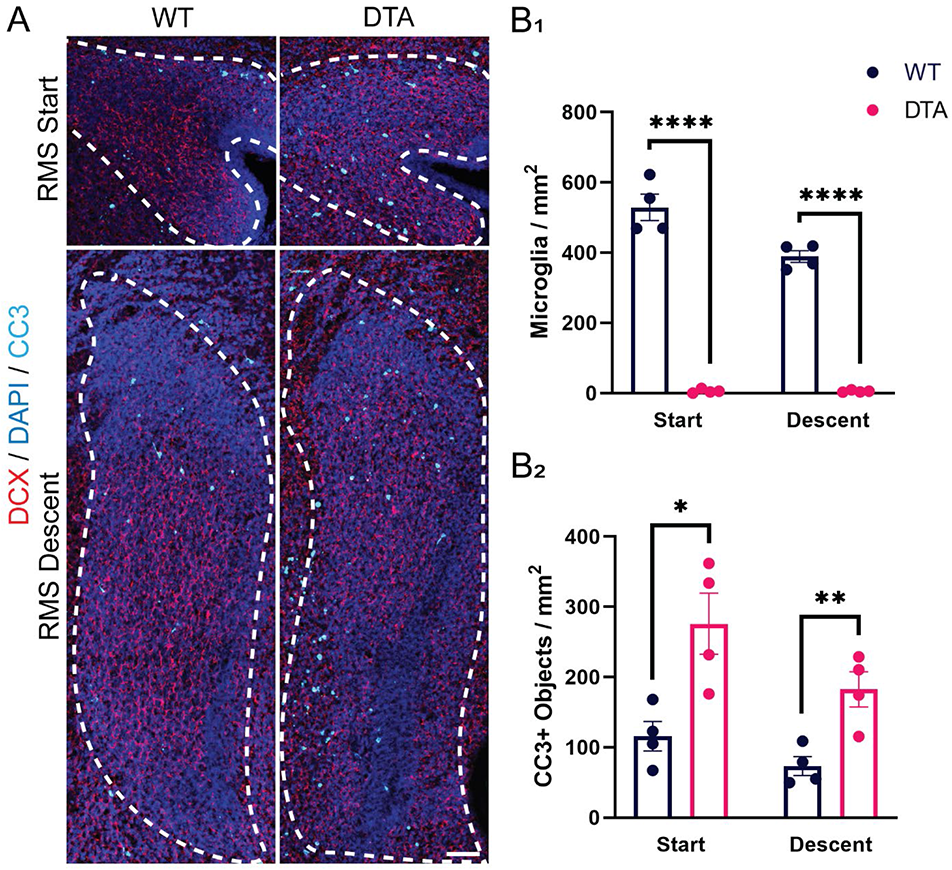
Microglia depletion for three days postnatally is associated with an accumulation of apoptotic cells in the rostral migratory stream (RMS). (A) Coronal sections of the start and descent of the RMS in *Cx3cr1^CreER/+^; ROSA26^+/+^* (left) and in *Cx3cr1^CreER/+^; ROSA26^eGFP-DTA/+^* (right) littermates following i.p. injection of tamoxifen at postnatal day 0 (P0) and sacrifice at P3. The RMS region was defined by increased density of DAPI nuclear staining, as shown by the white dashed outline. Scale bar = 50 μm. (B_1_) There was a significant decrease in microglia density following 3 days after tamoxifen injection in P0 *Cx3cr1^CreER/+^; ROSA26^eGFP-DTA^* mice. (B_2_) There was a significant increase in the density of CC3+ cells in the RMS of DTA+ mice following injection of tamoxifen in both the start of the RMS and RMS descent. *p <0.05; **p < 0.01; ****p<0.0001.

**Figure 12.**
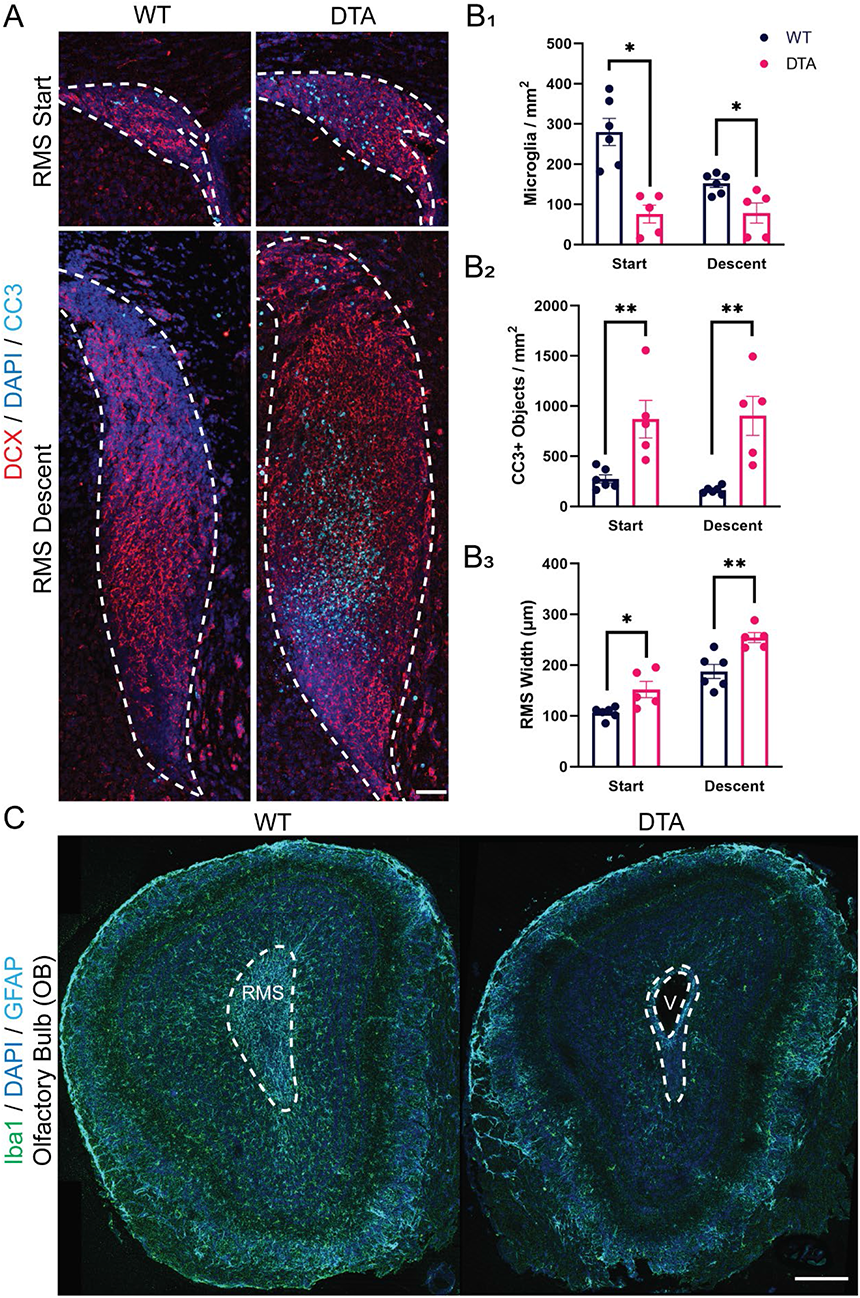
Microglia depletion for 14 days postnatally is associated with an accumulation of apoptotic cells and patent olfactory ventricles in the rostral migratory stream (RMS). (A) Coronal sections of the start and descent of the RMS in *Cx3cr1^CreER/+^; ROSA26^+/+^* (left) and in *Cx3cr1^CreER/+^; ROSA26^eGFP-DTA/+^* (right) littermates following i.p. injection of tamoxifen at postnatal day 0 (P0) and every three days subsequently until sacrifice at P14. Apoptotic cells were identified with cleaved caspase-3 (CC3) labeling (cyan) and neuroblasts with doublecortin (DCX) staining (red). Scale bar = 50 μm. (B1) There was a significant decrease in microglia density following 14 days of tamoxifen injection in *Cx3cr1^CreER/+^; ROSA26^eGFP-DTA^* mice. (B_2_) There was a significant increase in the density of CC3+ cells in the RMS of DTA mice following 14 days of microglia depletion in both the start of the RMS (WT vs DTA p = 0.0079) and RMS descent. (B_3_) There was also a significant increase in the width of the RMS at its widest point at both the RMS start and descent. The RMS region was defined by increased density of DAPI nuclear staining, as shown by the white dashed outline in A. (C) Coronal sections of the whole olfactory bulb (OB) in *Cx3cr1^CreER/+^; ROSA26^+/+^* (left) and in *Cx3cr1^CreER/+^; ROSA26^eGFP-DTA/+^* (right) littermates. Pups were injected every three days starting at P1 and sacrificed at P14. The RMS region is defined by increased density of DAPI nuclear staining in the OB core, as shown by the white dashed outline. A decrease in the number of microglia (green) is seen in the OB of the DTA+ mouse (right) following TMX injections. Enlarged and patent olfactory ventricles were observed in DTA+ mice (V), as opposed to the closed olfactory ventricles of WT littermates (left). Scale bar = 250 μm. *p <0.05; **p < 0.01.

Of further interest, we also found patent olfactory ventricles in the core of the RMS in all five mice that expressed the DTA+ subunit following 14 days of microglia depletion (Fig. 12C). This parallels early findings of enlarged and patent olfactory ventricles observed in three week old Csf1 null mice that have a significant depletion of microglia (Erblich et al., 2011). The enlarged olfactory ventricles may reflect increased ventricular pressure. It is possible that phagocytic microglia lining the RMS are responsible for clearing apoptotic debris, and thus an accumulation of debris in their absence may obstruct normal CSF outflow and cause ventriculomegaly.

## DISCUSSION

We sought to determine if the ameboid microglia that invade the RMS early in development play a role in facilitating neuroblast migration. We found that microglia line the outer borders of the RMS and intimately associate with migrating neuroblasts in the early postnatal period. Microglia ablation did not impact the migratory capacity of neuroblasts in the postnatal RMS. However, microglia express phagocytic markers and envelop migrating neuroblasts, implicating active phagocytosis of neuroblasts in the RMS. Furthermore, microglia depletion induced an accumulation of apoptotic and densely packed neuroblasts, an expanded RMS, and patent olfactory ventricles. Together, these results suggest that microglia function to maintain an environment permissive to neuroblast migration by clearing excess and possibly defective neuroblasts.

Microglia are present in the RMS during the first postnatal week, where they exhibit an ameboid morphology, accumulate at the borders of the RMS and closely associate with migrating neuroblasts. This microglia distribution occurs prior to the assembly of the vascular scaffold and astrocyte tube. There was only sparse CD31 blood vessel labeling in the RMS during the first postnatal week, which lengthened at P14 to resemble the scaffold that parallels the path of neuroblast migration in adults (Bozoyan et al., 2012). An astrocyte tube was similarly not observed during the first postnatal week. Instead, GFAP-stained radial glia radiated from the RMS core (Alves et al., 2002). However, neuroblast migration was orthogonal to these processes, suggesting that migration may be independent of radial glia (Alves et al., 2002). A fine network of astrocyte processes coalesced into thick septate processes at P21 that further consolidated into the densely packed astrocytic tube typical of adults at P56 (Peretto et al., 2005). The distribution of microglia flanking the outer borders of the RMS during the first postnatal week suggests that they may support the developing RMS in the absence of the vascular scaffold and astrocyte tube.

There is a preponderance of ameboid microglia in the RMS during the first postnatal week (Xavier et al., 2015) before they shift to a more ramified morphology, indicating that microglia functions may differ between these periods. “Dense” and ameboid microglia in the RMS decreased in number by P14, with a concomitant increase in microglia process complexity by P21. This change in microglia morphology from ameboid to ramified during development could simply be a manifestation of normal microglial cell development. Microglial cell development follows a step-wise developmental program with distinct temporal stages in gene expression (Matcovitch-Natan et al., 2016; Hammond et al., 2019; Li et al., 2019; Masuda et al., 2019, 2020). The transcription factors PU.1 and RUNX1 drive the transition from progenitor cells to microglia (Kierdorf et al., 2013). Transforming growth factor-beta 1 (TGF-β1), which drives PU.1 as well as the transcription factors MAFB, SALL1, MEF2A and EGFR1, is also necessary for proper cell development (Butovsky et al., 2014). Mutations in SALL1 induce an abnormal ameboid microglia morphology and downregulation of the homeostatic genetic signature (Buttgereit et al., 2016; Utz et al., 2020). Throughout microglial cell development there is a gradual expression of adult homeostatic genes, such as Tmem119, P2ry12, and Hexb (Bennett et al., 2016; Matcovitch-Natan et al., 2016).

Alternatively, varying microglia morphologies and shifts in gene expression could reflect different functional activities of microglia at different developmental stages. While single-cell RNA-sequencing revealed little microglia transcriptomic heterogeneity in adult mice, microglia isolated from developing mice exhibited heterogeneous gene expression profiles, suggesting diverse functionality (Hammond et al., 2019; Li et al., 2019; Masuda et al., 2019). Postnatal microglia transcript clusters further illustrated varied distribution across brain locations, implicating local specification of microglia function during development (Masuda et al., 2019). Meanwhile, whole genome transcriptional profiling of microglia showed enrichment of biologic processes related to neurogenesis and neural development during embryonic life (E14.5) and early postnatal life (P1), but not adulthood (Grassivaro et al., 2020). Thus, specific microglia subsets may occur in a spatially and temporally restricted matter to regulate specific neurodevelopmental events.

The intimate association between ameboid microglia and migrating neuroblasts in the early postnatal period, prior to the formation of the vascular scaffold and astrocyte tube, led us to consider the possibility that microglia facilitate neuroblast migration. However, microglia depletion for three days postnatally with two different ablation strategies did not alter the density of neuroblasts within the RMS nor the distance they migrated. These results indicate that microglia are not necessary for neuroblast migration in the early postnatal RMS. This is consistent with the findings of Kyle et al. (2019), who found no difference in the density of BrdU pulse labeled neuroblasts in the RMS of adult animals treated with the oral CSF-1R inhibitor PLX5622. In contrast, significantly fewer BrdU+ cells reached the glomerular layer (GL) in the OB of adult animals injected with a monoclonal saporin toxin conjugated to a CD11b monoclonal antibody to deplete microglia (Ribeiro Xavier et al., 2015). The accumulation of pyknotic apoptotic neuroblasts in the RMS following microglia depletion in this study may have constrained the ability of neuroblasts to reach the OB GL. Alternatively, microglia may promote neuroblast radial migration in the OB, as opposed to the tangential migration in the RMS. Microglia secrete chemokines known to be involved in neuroblast radial migration, including IGF-1 (Suh et al., 2013; Ueno et al., 2013) and PROK2 (Mundim et al., 2019). We also saw a decrease in the density of BrdU+ cells in the OB GL after microglia depletion. Thus, while microglia depletion does not impact the migratory capacity of neuroblasts in the RMS, it does appear to impede the ability of neuroblasts to reach their ultimate destination in the OB.

Microglia may be essential for maintaining an environment permissive to neuroblast migration in the RMS through phagocytosis. Microglia lining the RMS in the first postnatal week express the phagocytic markers CLEC7A, MERTK and CD68 (Fourgeaud et al., 2016; Li et al., 2019b); in contrast, microglia in the cortical regions outside the RMS do not, implicating a specific role for microglia phagocytosis in the RMS. Microglia expressing phagocytic markers further “wrap” neuroblasts during this period in the RMS, suggesting phagocytosis of actively migrating neuroblasts. Microglia also phagocytose labeled antigens in distal regions of the RMS, indicating the monitoring and phagocytosis of extracellular material. The broader domain of the RMS packed with CC3+ neuroblasts after 14 days of postnatal microglia depletion demonstrates an accumulation of apoptotic or defective neuroblasts. Furthermore, improper clearance of antigens and apoptotic debris in the RMS could induce increased CSF pressure and explain the enlarged and patent olfactory ventricles following microglia depletion (Erblich et al., 2011; Hsu et al., 2019). An accumulation of neuroblasts and apoptotic debris in the RMS may similarly impede neuroblast migration over time. Together, these findings illustrate the importance of microglia phagocytosis in sustaining RMS homeostasis.

The accumulation of neuroblasts within a wider RMS following microglia depletion indicates that a surplus of neuroblasts are generated during development, which are subsequently eliminated by microglia. However, it remains unexamined whether microglia simply phagocytose dead or dying neuroblasts in the RMS, or if they “kill” microglia in the RMS via phagoptosis. There is evidence that microglia can “kill” cells in neurogenic regions through an extracellular respiratory burst (Marín-Teva et al., 2004; Sierra et al., 2010). Microglia depletion caused an increase in Purkinje cell number in the cerebellum, suggesting that microglia eliminated “living” and not apoptotic Purkinje cells (Marín-Teva et al., 2004). Microglia also phagocytosed mitotically active neural precursor cells independent of apoptotic pathways in the SVZ (Cunningham et al., 2013). Furthermore, despite the accumulation of apoptotic cells in the RMS of *Axl* and Mertk knockout animals, there was a ∼70% increase in the cellular density of the granule cell and glomerular layers of the OB, suggesting that many neuroblasts phagocytosed by microglia are not apoptotic but instead ‘eaten-alive’ (Fourgeaud et al., 2016). Future studies may reveal whether microglia randomly target migrating neuroblasts for phagoptosis or are able to detect aberrantly migrating or otherwise defective neuroblasts.

An alternative explanation for the increase in apoptotic neuroblasts following microglia depletion in the RMS is that microglia provide trophic support for neuroblasts. Microglia depletion caused an accumulation of BrdU+ fragments within pyknotic neuroblast nuclei in the adult RMS (Ribeiro Xavier et al., 2015). There may be regional heterogeneity of microglia functions within the RMS, with some microglia being responsible for trophic support and others for neuroblast elimination. Such heterogeneity was demonstrated in ameboid microglia expressing SPP1 in the corpus callosum during the first postnatal week: microglia expressing high transcript levels of *Spp1* neighbored microglia with no expression (Hammond et al., 2019). Paralleling this finding, SPP1 was also observed in a subset of CLEC7A+ microglia in the developing corpus callosum of P7 adjacent to CLEC7A+ with no SPP1 expression (Li et al., 2019a). Another candidate trophic factor that may support neuroblasts during the postnatal period is IGF-1, which is expressed by the CD68+ CLEC7A+ microglia subtype (Hammond et al., 2019; Li et al., 2019a) and supports oligodendrocyte precursor cells (Wlodarczyk et al., 2017) and projecting axons (Ueno et al., 2013) in the first postnatal week. There may therefore be local heterogeneity of microglia functions in the early postnatal RMS, with microglia responsible for neuroblast elimination neighboring microglia that support neuroblast survival.

In summary, microglia phagocytosis is crucial for maintaining the homeostatic conditions in the RMS that allows for effective neuroblast migration. A role of microglia for clearing defective cells via phagocytosis may be reflected in the child born without microglia (Oosterhof et al., 2019); the observed heterotopias may be a result of a lack of appropriate phagocytic clearance of defective neuroblasts, as opposed to a general failure of neuroblast migration in the absence of microglia. Microglia may be able to identify cells with chromosomal errors or that exhibit oxidative stress (Cunningham et al., 2013). Alternatively, microglia may simply regulate the size of the neuroblast pool and eliminate cells that would otherwise be viable. Phagocytosis of excess or defective neuroblasts and clearance of apoptotic cells and debris likely enables an environment permissive to neuroblast migration in the RMS. As our understanding of the functional capabilities of microglia and their importance in regulating neurodevelopment continues to expand, these results shed important insight into the role of microglia in maintaining the homeostasis of neuroblast migratory corridors during the early postnatal period.

## Acknowledgments

We thank Christine Kaliszewski for assistance.

